# Activating NK- receptors, homing selectins and inhibitory Siglecs recognize EBOLA-GP and HPV-L1NK

**DOI:** 10.1101/2020.07.24.219329

**Authors:** Mostafa Jarahian, Katharina Marstaller, Heribert Wurmbäck, Nadine Banna, Roshanak Ahani, Hossein Etemadzadeh, Lea Katharina Boller, Kayhan Azadmanesh, Angel Cid-Arregui, Martin R Berger, Frank Momburg, Carsten Watzl

**Affiliations:** Toxicology and Chemotherapy Unit, German Cancer Research Center (DKFZ), Heidelberg; Department of Virology, Pasteur Institute of Iran; Targeted Tumor Vaccines in German Cancer Research Center (DKFZ), Heidelberg; Antigen Presentation and T/NK Cell Activation, German Cancer Research Center (DKFZ), Heidelberg; Department of Immunology, Leibniz-Research Centre for Working Environment and Human Factors at the TU Dortmund (IfA-Do), Dortmund, Germany

**Keywords:** Siglecs, Ebola virus glycoprotein, HPV, NCRs, NKp44, NKp46, selectins

## Abstract

The Ebola virus glycoprotein (EBOV)-GP is extensively glycosylated. Its expression induces a physical alteration of surface adhesion molecules, which causes cell rounding and detachment of the infected cells. This phenomenon likely plays a crucial role in viral pathogenicity. In this study, we show that such morphological changes are cell line-dependent as well as dependent on the surface proteins that interact with EBOV-GP in *cis* and *trans*. We have generated data showing that natural killer (NK) cell receptors (NCRs: NKp44 and NKp46), selectins (CD62E/P/L) and inhibitory Siglecs function as receptors for Ebola-GP and human papilloma virus (HPV-L1). We used HEK293 cells transfected with Ebola-GP and recombinant fusion proteins containing the extracellular domain of each of these receptors linked to the Fc of human IgG1, which showed significant differences in their virus-binding behavior compared to HEK293 cells transfected with empty vector. Further, to demonstrate that EBOV-GP is a ligand for NKp44 and other NK-receptors, and to investigate their role in immune escape, we also used human HEK-293, HeLa- and hamster CHO-GP-transfectants. Our data show that the NK receptors NKp44 and NKp46 play a key role in recognizing EBOV (Ebolavirus) and strongly suggest that other inhibitory (Siglec-7, Siglec-5) and non-inhibitory homing receptors (P-Selectin, L-Selectin, E-Selectin, and DC-SIGNR/DC-SIGN) take part in the interaction with virus particles. In addition, we show that NKp44, and NKp46, Siglec-7, and -5, and P-, L-, E-selectins as well as of and DC-SIGNR/DC-SIGN bind to the artificial viral envelope of a lentiviral vector that contains EBOV-GP. Altogether we prove that NCRs and a range of other inhibitory and activating receptors can interact with viral envelope/capsid proteins and that such interaction could play an important role in the elimination of virus infected cells. Our findings could be used to develop new strategies for prevention and treatment of infections by these viruses.

**Author summary:** The innate immune system is able to recognize specifically certain virus components. Here we show that activating NK-cell receptors (NKp44, and NKp46) are involved in such interaction by using HEK293 and CHOK1 cells transfected with the Ebola virus glycoprotein (EBOV-GP) and by binding studies with purified EBOV-GP. In detail, we have found moderate to strong affinity of Siglecs (Siglec-7, and -5), selectins (P-, L-, E-Selectin) and DC-SIGNR/DC-SIGN to purified EBOV-GP, and to cells transfected with EBOV-GP as well as to the envelope of a lentiviral vector carrying the EBOV-GP. Our findings show that NKp44, and NKp46, Siglec-7, and -5, as well as P-and L-selectins have a strong affinity to EBOV-G.

## Introduction

Ebola- and Marburgvirus are two genera of the Filoviridae family belonging to the most virulent viruses known, which in humans cause a rapidly fatal hemorrhagic fever (1, 2). Ebolavirus has caught worldwide attention, when in 2014 an epidemic spread across West Africa caused hemorrhagic fever and the death of many (2–6). Ebolavirus is able to infect almost every cell type, with a rapid rate of viral replication (7). Treatment is still symptomatic, as an effective virostatic agent has not been approved yet.

Ebolavirus contains seven different genes (3’-NP-VP35-VP40-GP-VP30-VP24-L-5’), out of which the GP gene forms nine different glycoproteins (GPs) via alternative open reading frames. The other six genes code for structural proteins of Filovirus particles. These GPs can be modified in different ways by various enzymes. After N- and O-glycosylation, GP0 is cleaved by the enzyme furin to yield the GP1 and GP2 subunits (8–10). GP is structured in a chalice-like shape with a trimer of GP1 / GP2 heterodimers, out of which GP2 is forming the base and GP1 the cup (11). This GP-trimer can be cleaved from the viral surface via the tumor necrosis factor-α-converting-enzyme (TACE) (11). GP1 is mucin-like and contains N-glycosylated-as well as O-glycosylated areas, where mutations permanently take place, thus enabling immune escape (12, 13), whereas GP2 has only two N-glycosylated areas (14, 15). In addition to the membrane-bound proteins GP1 and GP2, the Ebola virus (EBOV) secretes multiple soluble glycoproteins (sGP), as a result of differential splicing of the glycoprotein gene transcripts (10, 11, 16–18). The entry of filovirus is mediated by the viral spike glycoprotein (GP), which attaches to viral particles on the cell surface and allows fusion between endosomes and the virus particles on the cell membrane (19, 20). These proteins, despite their large common primary sequence, differ significantly in their structure. Soluble glycoproteins share the 295 N-terminal amino acids with GP1, but have a unique carboxyl terminus of 28 amino acids (16). Various in vitro studies addressing EBOV-GP presence on the cell surface suggest that its expression impairs the cell-cell interaction and facilitates cell rounding and detachment. Furthermore, detachment-mediated apoptosis was observed in endothelial cells expressing EBOV-GP (21–24). GP1 is responsible for receptor binding (including α-dystroglycan, heparan sulfate, DC-SIGN, etc.) and GP2 mediates low pH-induced membrane fusion. Both proteins interact with each other to form a stable homotrimer complex on the viral envelope (25, 26).

Currently, the only known function of sGP is its anti-inflammatory effect on endothelial cells treated with TNF-α, an effect that was thought to interfere with the recruitment or extravasation of leukocytes (10, 27–29). sGP is post-translationally modified by N-glycosylation, but also by C-mannosylation (9). Fuller and colleagues have shown that both, Marburg virus-like particles (mVLPs) and Ebolavirus-like particles (eVLPs), trigger a consistent upregulation of CD69 on the cell surface of polyclonal NK cells from different donors (30). Upregulation of CD69 on NK cells is closely linked with activation of several signal transduction pathways including survival and induction of cytokine production and cytolysis of targets (31, 32). Fuller and colleagues demonstrated a correlation between increased expression of sialic acid on the surface of infected cells with resistance against activation of the complement system (30). EBOV infection is able to impair type-I IFN production by infected cells and block IFN response in uninfected cells (33, 34). In addition, EBOV infection is able to induce massive NK cell apoptosis, thus avoiding NK function and impairing NK-mediated DC maturation (35–38).

The killer cell immunoglobulin-like receptors (KIRs) as well as Siglecs, and CD94-NKG2A, are involved in the inhibitory signal cascade of NK cells. Most inhibitory receptors recognize specific MHC class I isoforms and thereby ensure tolerance of NK cells against self-antigens (39). NK-cell-activation involves KIR2DS1-5, NKG2D, CD16, and NCRs (natural cytotoxicity receptors NKp46, NKp44 and NKp30). Treatment with mAbs against NCRs results in a very effective block of NK cell activity and thus reduces NK cell cytotoxicity against certain tumor cells (40, 41).

NKp46 (NCR1) and NKp44 are expressed by innate lymphoid cells (ILCs) of group 1 (ILC1), a subset of group three ILCs (NCR + ILC3) (42), and by γδ T cells (43). NKp46 is a 46 kDa type I transmembrane protein belonging to the immunoglobulin (Ig) superfamily, which is characterized by two extracellular C2-type Ig-like domains (D1 and D2) (40, 41) (Fig. 1). The TM domain of NKp46 contains a positively charged arginine residue that mediates association with the negatively charged aspartate residue found in the TM domain of the ITAM signaling adaptors, CD3ζ or the Fc receptor common γ (FcRγ) (41) (Fig. 1). NKp44 (NCR2) is a 44 kDa protein, consisting of a single extracellular V-type Ig (IgV) domain (44–46), followed by a long stalk region and a hydrophobic transmembrane domain. The latter contains a charged lysine residue that leads to association with DAP12 adaptor with ITAM (47) (Fig. 1). Structural studies have shown that the IgV domain of NKp44 forms a saddle-shaped dimer with a positively charged groove on one side of the protein (47, 48). The NKp44-1 isoform has potential inhibitory ITIM but also the capacity to perform an activating function via association with DAP12 (49).

**Fig. 1.**
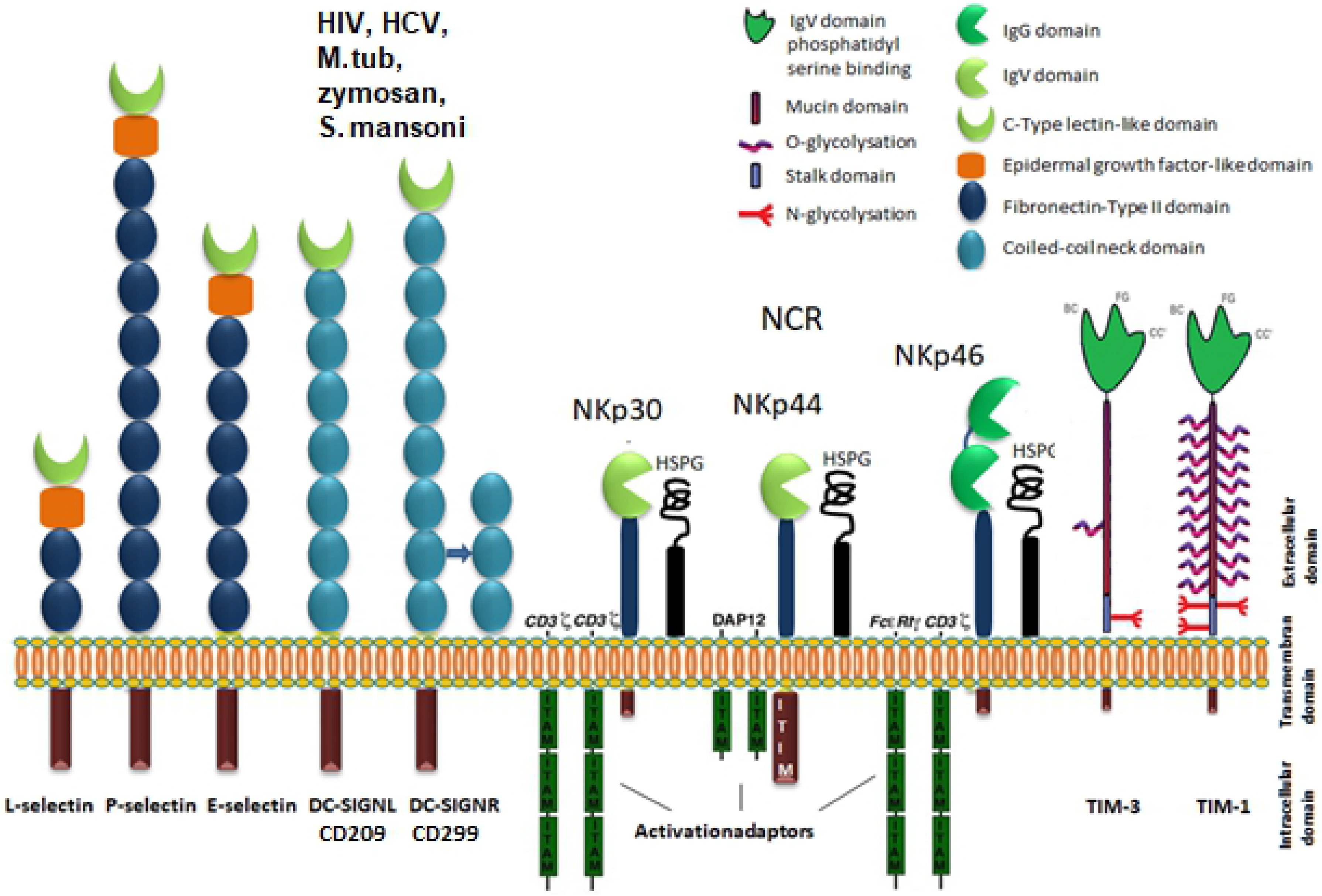
Activating and inhibitory receptors binding EBOV-GP, which are expressed on the surface of immune and non-immune cells. From left to right: L, P and E selectins on lymphocytes, platelets and epithelial cells, respectively, DC-SIGN (Dendritic Cell-Specific adhesion molecule), DC-SIGNR on NK cells, and the human natural cytotoxicity receptors (NCR) can recognize glycan epitopes of EBOV-GP (envelope glycoprotein GP) during immune responses against infected cells. The Tim family members (TIM-3, TIM-1) recognize phosphatidylserine (PtdSer), a component of the Ebola virus envelop that promotes the cell entry of virus, but not GP.

NKp30 (NCR3) is a 30 kDa protein that, similarly to the NKp46, is expressed on all mature resting and activated NK cells as well as on ILC2 cells (50). It contains an IgV domain and a hydrophobic transmembrane domain with a charged arginine residue. This leads to its ability to associate with the ITAM adaptors, CD3ζ and/or FcRγ (51, 52) (Fig. 1). There are six alternatively spliced transcripts from the NKp30 gene, termed NKp30a-f. As a consequence, NKp30 and NKp46 surface expression on adaptive memory NK cells is reduced (53). Cellular ligands for NKp30 include nuclear factor BAT3 and the he B7 family B7-H6 (54, 55). In previous work, we were able to show that ligands for NKp30 and NKp44 can be detected on the surface and in intracellular compartments of different tumor cells (56). These ligands often contain heparan sulfate linked to proteoglycans (57, 58). Further we reported that NKp30 and NKp46 present on natural killer cells play a key role in the immune response in the presence of vaccinia and mouse pox virus (ECTV/ectromelia virus), as they bind to hemagglutinin (HA)- a component of the Vaccinia virus envelope (59). It should be noted that NKp30 triggered activation of NK cells is blocked by HA of Vaccinia virus, whereas NKp46 stimulates NK cells (59). Moreover, the pp65 matrix protein of human cytomegalovirus (HCMV) binds NKp30 and inhibits its function (60). The results indicate that NKp30 has a different role in NK-cell cytotoxicity (51, 59, 61). Further, interferon (IFN)-γ secretion by NK and non-NK cells plays a role in the antiviral effect (62, 63). For example, type I interferons are essential for the activation of NK cells during vaccinia virus (VV) infection (64). A VV infection induces expression of ligands for the activating natural cytotoxicity receptors (NCRs), and increases the susceptibility of host cells to lysis by NK cells (65). In addition to VV, human immunodeficiency virus and herpes simplex virus also contribute to an upregulation of the expression of NCR ligands on infected cells (66, 67).

NKp46 recognizes the sigma1 protein of reovirus (68). In addition, the HA proteins of sendai-, influenza-, and newcastle disease (ND) viruses are able to bind NKp46 and NKp44, and induce NK cell activation (69–72). In the human influenza and ND viruses, this is achieved when HA on the virus capsid links to sialic acid residues located on the surface of target cells. For cellular uptake, another structural viral protein, termed neuraminidase, cleaves the glycosidic bond to sialic acid residues, and thus liberates the virus for fusion with the target cell membrane (59, 69, 71–73). NKp44 interacts with envelope glycoproteins from the West Nile and dengue viruses E/M proteins (74).

DC-SIGN (CD209) and DC-SIGNR (DC-SIGN-related, CD299, CLEC4M) bind soluble ebola- glycoproteins (EBOV-GP) with similar affinity (75), as well as soluble human immunodeficiency virus type-1 (HIV-1) gp120. This interaction is inhibited in an environment with increased pH (75). Highly glycosylated gp120 on the HIV-1 envelope has strong binding avidity to tetramerized DC-SIGN clusters on the surface membrane of dendritic cells. DC-SIGN and DC-SIGNR are furthermore receptors for other virus proteins such as from Lassa and Hepatitis C viruses (76–78).

DC-SIGN and DC-SIGNR are calcium-dependent C-type lectins, which have high affinity for ICAM3 (CD50) (79). DC-SIGN has low affinity to weakly polysialylated NCAM-1 (80). Lysis of dendritic cells by NK cells with polysialylated CD56^dim^ is increased in the presence of an anti-DC-SIGN antibody, which inhibits the interaction between DC-SIGN and its ligands (e.g. NCAM-1) (80). Several unrelated viruses, including influenza virus, RSV and HIV can directly suppress NK cell activity. Binding to DC-SIGN can promote HIV and hepatitis C viruses to infect T-cells via dendritic cells (77, 81, 82). HIV-infected lymphocytes (viral core protein p24 positive) do not present detectable DC-SIGN ligands (ICAM-1, ICAM-3, DC-SIGNR) on their cell surfaces, but non-infected lymphocytes, carrying NK-specific marker NCAM1, present cell surface DC-SIGN ligands (ICAM-1, ICAM-3) (80). HIV infection causes leukocyte dysfunction e.g. by increasing the level of soluble adhesion molecules (ICAM-1, soluble L- and E-selectins) leading to altered cell interactions (83).

L-selectin is a type I transmembrane cell adhesion molecule expressed on most circulating cells, including neutrophils, dendritic cells, monocytes, B cells, NK and T cells. L-selectin is a major regulator for leukocytes during trans-endothelial migration. E- and P-selectin are expressed on endothelial cells at sites of inflammation, interact with receptors on the surfaces of leukocytes, while L-selectin is expressed on lymphocytes and binds to glycans on high endothelial venules of lymph nodes. The glycoprotein gp120 of HIV-1 binds L-selectin in solution and on the host cell membrane. Upon entry of HIV into CD4^+^ T cells, L-selectin is cleaved at the membrane proximal site by proteolysis, thus facilitating virus release from cells (84, 85). L-selectin is clustered with E-selectin and CD34 in trans (86, 87), whereby co-localization of L-selectin with ADAM17 is induced to the uropod (85).

Recently it was found that TIM-1 (T-cell-immunoglobin and mucine-domain 1) is a filovirus-receptor (88) and interacts by its PtdSer binding pocket directly with phosphatidylserine (PtdSer) located on the viral capsid (89–91). Furthermore, TIM-1 binds the adhesion receptor P-selectin and mediates T cell trafficking during inflammation and autoimmunity (92, 93). TIM-1 can regulate and enhance type 1 immune response against tumors (94). Filoviruses attach to the cell membrane via non-canonical cell surface receptors, C-type lectins (CLECs), and PtdSer receptors. CLECs (DC/L-SIGN, LSECtin, ASGPRI, and hMGL) interact with sugars on the virion’s glycoprotein through their carbohydrate recognition domains. TAM family members are also PtdSer receptors and interact with PtdSer binding proteins Gas6 or Protein S (95–98).

An effective immune response must result from coordination between the activities of the humoral and cellular immune systems. EBOV sGP and glycosylated-GP (GlycGP) modulate host immune responses (99). The EBOV-infection of DCs leads to deregulation of signal amplifications and polarization between DC and T-effector cells, which takes place between MHC-I and MHC-II peptide presentations and interaction with the TCRs on T-cells. These inappropriate DC/T-cell interactions can lead to apoptosis in T-cells, which will eliminate clonal expansion of CD4 helper T cells, block T-CD8 mediated cytotoxicity and antibody production by B cells.

## Results

### EBOV-GP binds NCRs, homing selectins and inhibitory Siglecs

We used purified EBOV-GP coated ELISA-plates to assess the binding to IgG1-Fc recombinant fusion proteins carrying NKp44-Fc, CD44-Fc, CD24-Fc, PSGL-1-Fc and the known Ebola receptors DC-SIGN/DC-SIGNR. Our results show differential binding of EBOLA-GP to these receptors. DC-SIGN (CD209) and its related C-type lectin DC-SIGNR (CD299) bind strongly to EBOV-GP, as previously shown (75). NKp44 binds moderately to EBOV-GP. NKp30-Fc, CD44-FC, PSLG-1-Fc, NKG2D-Fc and CD24 shed from placenta have no significant binding to the EBOV-GP (supplementary Fig1). We next extended our study to test the binding of NKp46-Fc, P-selectin-Fc, E-selectin-Fc, L-selectin-Fc and inhibitory Siglec7-Fc, Siglec5-Fc, Siglec3-Fc. In order to reduce non-specific binding of some fusion proteins with high binding capacity (e.g. P-selectin-Fc, Siglec5-Fc) we tested different protein-containing and protein-free blocking buffers. The results show that P-selectin-Fc, L-selectin-Fc (CD62P/L) and inhibitory receptor Siglec3, Siglec5-Fc, Siglec7, as well as DC-SIGN/DC-SIGNR bind to EBOV-GP to varying degrees (Fig. 2). Out of these, Siglec3-Fc and E-selectin-Fc showed relatively weak binding to EBOV-GP, while NKp30-Fc, PSGL-Fc, CD44-Fc, and CD24-Fc do not bind significantly to EBOV-GP, compared to the controls. These results were confirmed repeatedly under different conditions and with different batches of proteins and the binding tendencies were reproducible throughout all experiments (Fig. 2).

**Fig. 2:**
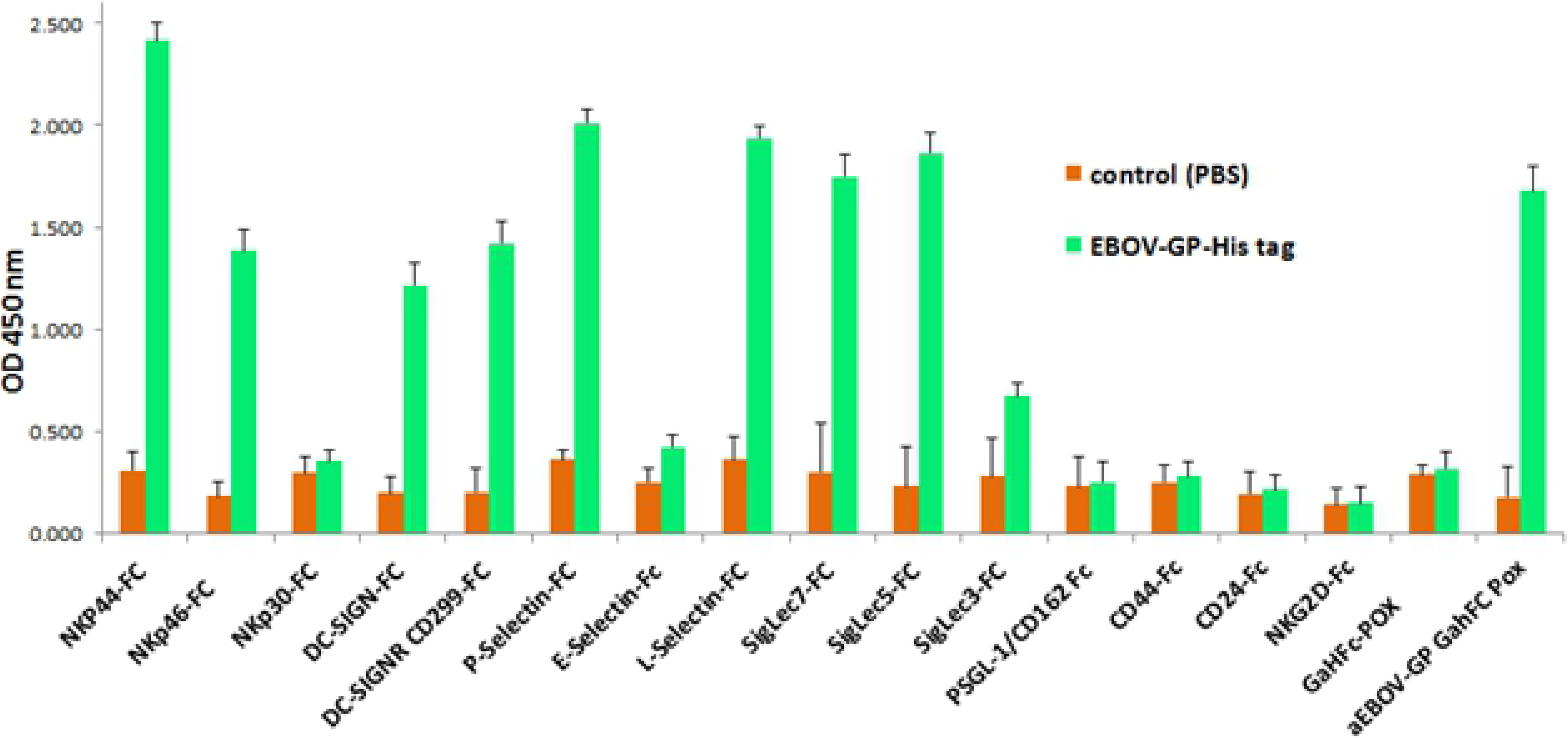
Binding of EBOV-GP-His-tag to activating receptors, homing proteins (selectins) and inhibitory siglecs. Purified EBOLA-GP-His-Tag coated ELISA-plates were used to test the binding of the indicated recombinant proteins. GaHFc-POD and GaMFc-POD served as negative controls and haEBOLA-GP as positive control. NCRs (NKp44, NKp46), L-Selectin, P-Selectin (CD62e/l/p), inhibitory receptor siglecs (siglecs-5, siglecs-7), and DC-SIGNR/DC-SIGN bind to EBOV-GP to varying degrees. From these, E-Selectin, and Siglec3 form a relatively weak bond to EBOV-GP compared to the control, while NKp44, NKp46, P-Selectin, L-Selectin, Siglec-5, DC-SIGNR, DC-SIGN and Siglec-7 bind significantly stronger to EBOV-GP than their controls. NKp30-Fc, PSGL-1-Fc, NKG2D-Fc, PSGL-Fc, CD44-Fc and CD24-Fc do not bind to the EBOV-GP in comparison to the respective controls. Furthermore, P-Selectin and Siglec-5 show a larger degree of unspecific binding compared to the other recombinant proteins.

### Expression of EBOV-GP in HEK-239 and CHO-K1 cells

EBOV-GP is highly glycosylated and interacts with various surface proteins simultaneously, thus affecting the adhesive behavior of infected cells. For instance, after transfection with EBOV-GP, HeLa cells change their morphology and behavior to grow as non-adherent cells at a slow proliferation rate (data not shown). We have transfected CHOK1 and HEK-239 cells stably with different plasmids carrying the EBOV-GP gene. It is known that HEK-293 cells are particularly sensitive to the expression of EBOV-GP viral glycoprotein (22, 24, 100). In order to investigate the binding behavior of normally glycosylated EBOV-GP, HEK-293 cells were separately transfected with two different expression-vectors, pCAGGS.cm5 (+)/GP (Marburg) and pcDNA6 (+)/GP-7916-5 (Heidelberg). CHOK1 was transfected with the expression-vector pCDNA 3.1. Both plasmids (pcDNA6 and pCDNA 3.1) contain a CMV (cytomegalovirus)-promoter, while pCAGGS-EBOV-GP consists of a chicken b-actin / rabbit β-globin hybrid. These two different species [human (HeLa, HEK-293) and Chinese hamster ovary (CHO)] were used, as we aimed to investigate the different patterns of glycosylation and *cis* co-partners for GP at the cell surface of transfected HEK-293 cells in human vs non-human cells and their respective effect on the interaction between EBOV-GP and the binding partners mentioned above. To determine GP-expression efficiency, we used a human monoclonal anti-ZEBOV GP antibody KZ52(101) with high affinity and a mouse monoclonal antibody (3B11) against EBOV-GP showing weak binding to GP-transfected HEK-293 cells. The latter anti-GP antibody showed a trypsin-dependent binding of EBOV-GP in HEK-293-cells, while the human anti-GP did not show dependency on trypsin (Fig. 3). The binding of 3B11 antibody (102) was twofold increased after trypsin treatment of GP-transfected cells (Fig.3a left). This may be due to the fact that EBOV-GP is present on the surface of transfected human HEK-293 cells in a highly glycosylated manner, leading to interaction with neighboring surface adhesion-proteins in *cis*. Apparently, expression of the EBOV glycoprotein results in correct processing and surface expression. The glycosylation pattern of the protein on the cell surface is broadly similar to the glycoprotein of the viral envelope. However, the binding analysis of EBOV-GP transfected HEK 293 cells showed different binding patterns with the fusion proteins, pending on the level of GP expression (Fig.4c). In addition, the fusion proteins have more than one cellular co-partner at the cell surface, which makes the comparison extremely difficult; hence, our strategy was to use a cell line from another species (e.g. CHOK1).

**Fig. 3:**
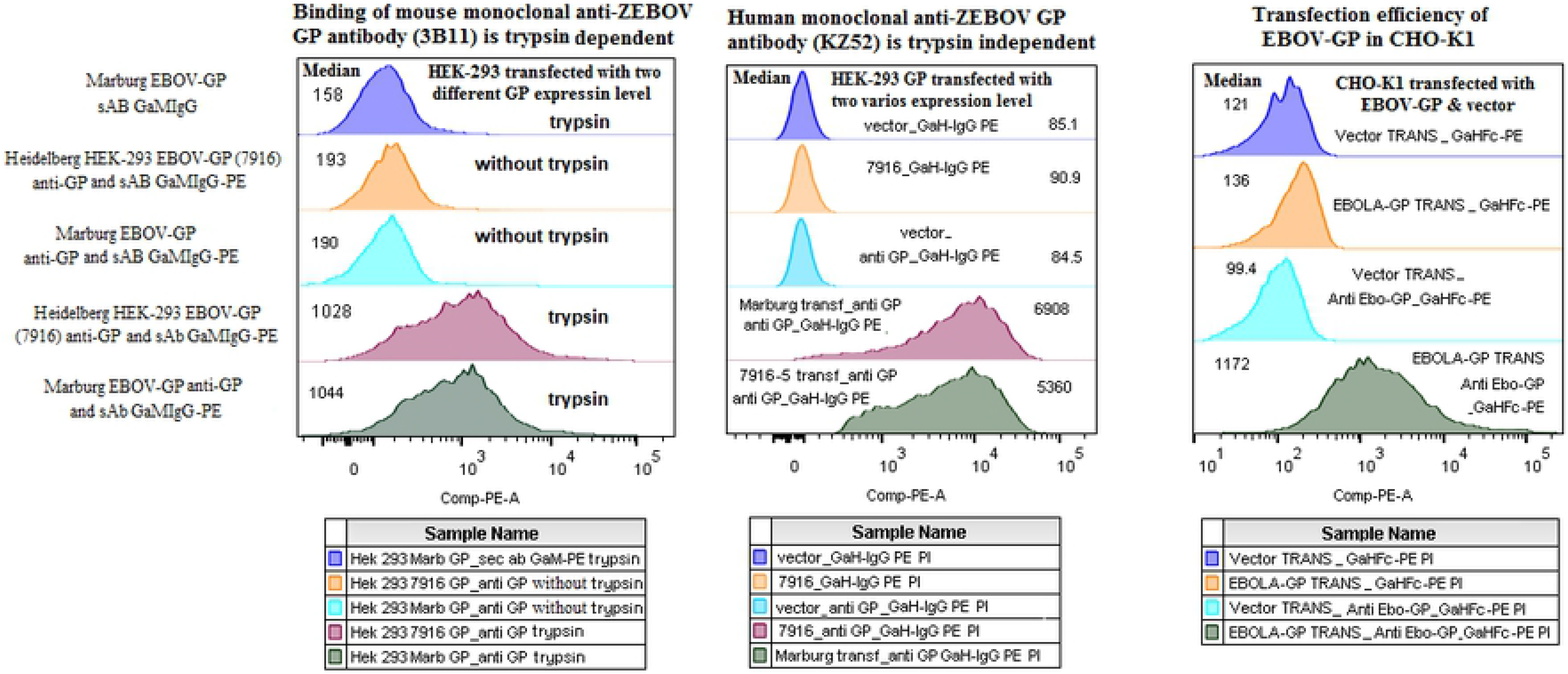
Expression efficiency of EBOV-GP in transfected CHOK1- and HEK-293 cells. Plasmids from Marburg (pCAGGS.cm5 + GP) and Heidelberg (pcDNA™6/V5-His + GP) were used for transfection of HEK-293 cells (3a), while the the plasmid pCDNA 3.1+ GP was used in CHO-K1 cells (3b). GP expression was detected by anti-ZEBOV GP mouse antibody (3B11), and anti-ZEBOV GP human antibody (KZ52). The binding to EBOV-GP transfected HEK-293-cells by the anti-GP mouse antibody 3B11 was trypsin-dependent (3a left). The human anti-GP did not show a dependency on trypsin (3a right). After transfection, the strongly positive cells were sorted and further checked for their degree of EBOV-GP expression by mAb anti EBOV-GP (KZ52).

**Fig. 4:**
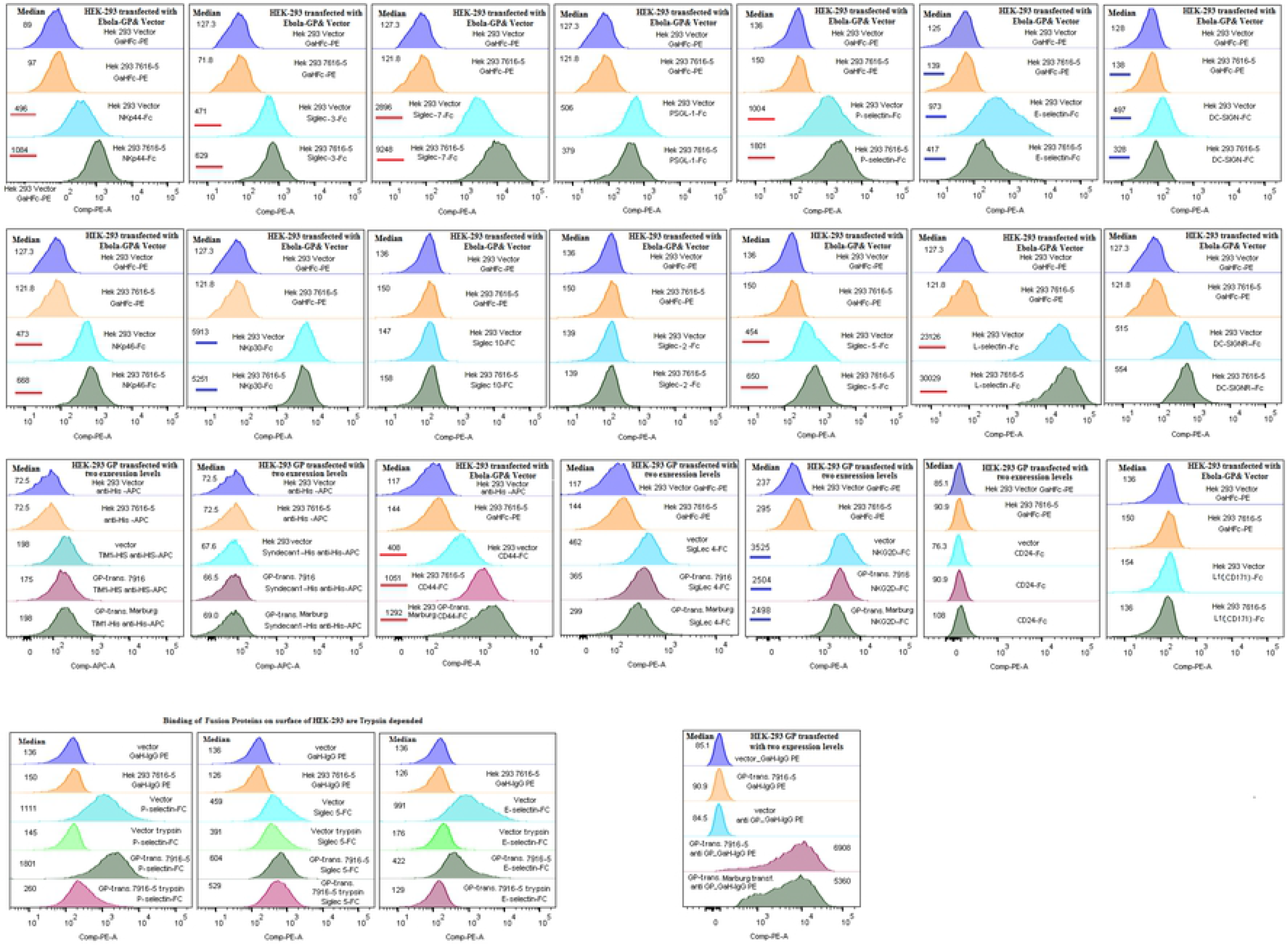
NCRs, homing selectins and inhibitory Siglecs bind to HEK293^EBOV-GP^ cells: 4a. Analysis of the binding of different fusion proteins to HEK-293^EBOV-GP^ cells. The figure shows that Siglec 7, NKp44, and L-selectin-Fc have a higher affinity to the EBOV-GP transfected cells, while Nkp46, Siglec-3, P-selectin, siglec-5-Fc and CD24-Fc have an only slightly increased affinity than the respective controls. NKp30-Fc, E-selectin-Fc, NKG2D-Fc, and CD44-Fc have a slightly reduced affinity to the EBOV-GP transfected cells. The binding of PSGL-1-Fc, siglec-2-Fc, siglecs-4-Fc, siglec10-Fc, TIM-1-Fc and DC-SIGNR/DC-SIGN does not change. 4b: Examples are shown for strongly trypsin-dependent binding of some recombinant fusion proteins (e.g. selectins) to their co-partner on the surface of target cells. 4c: Transfection efficiency of two different HEK-293^EBOV-GP^ cells.

### NCRs, homing selectins and inhibitory Siglecs bind to EBOV-GP transfected HEK-293 and CHO-K1 cells

Next, we used our fusion proteins (NCRs and the well-known Ebola-GP ligands CD209/CD299 (DC-SIGN/DC-SIGNR) for immunofluorescence staining of the human cell line HEK-293^EBOV-GP^. We observed strong, moderate or relatively week but significant binding of the fusion proteins to HEK-293^EBOV-GP^ cells in comparison to control vector-transfected HEK-293 cells (Fig.4a). We next tested the binding of the fusion proteins (NKp44-Fc, NKp46-Fc, NKp30-Fc, P-selectin-Fc, L-selectin-Fc, Siglecs2/ 3/ 4/ 5/ 7/ 10-Fc and CD44-Fc, CD24-Fc, PSGL-1-Fc, TIM1-Fc). Our results showed no significant binding of DC-SIGNR (CD299, DC-SIGN-related) to EBOV-GP transfected and control (vector) HEK-293 cells, and on the contrary, the binding of its co-partner DC-SIGN (CD209) was slightly decreased. The binding of L-Selectin-Fc, NKp44-FC, Siglec7-Fc to HEK-293^EBOV-GP^ cells was strongly enhanced, while NKp46, Siglec3-Fc, Siglec5-Fc, P-selectin-Fc and CD24-Fc showed moderate binding to HEK-293^EBOV-GP^ (Fig.4a). In comparison, there was no binding of NKp30-Fc, E-selectin-Fc, NKG2D-Fc, Siglec-4-Fc and DC-SIGNR (CD299)-Fc to HEK-293^EBOV-GP^ cells, which was at a level even lower than that to respective control transfected cells. The binding of DC-SIGN (CD209)-Fc, PSGL-1-Fc, Siglec-2-Fc, Siglec10-Fc, L1-CAM–Fc, and TIM-1-Fc did not change (Figure 4a). We could not prove binding of the fusion proteins CD44-Fc, CD24-Fc, PSGL-1-Fc and TIM1-Fc to purified EBOV-GP on ELISA plates. In contrast, the binding of proteoglycan CD44-Fc and glycoprotein CD24-Fc (selectin receptors) to HEK-293^EBOV-GP^ cells was significantly enhanced compared to the respective controls. In contrast, we could prove binding of the fusion proteins CD44-Fc, CD24-Fc, PSGL-1-Fc and TIM1-Fc to the artificial viral envelope of a lentiviral vector that contained the EBOV-GP, which was harvested from HEK-293^EBOV-GP^ cells (Fig.8). Interestingly, selectins, but not Siglecs showed reduced binding to HEK-293 cellular ligands upon pretreatment with trypsin, independent of the EBOV-GP presence (Fig. 4b).

### Binding of the fusion proteins to EBOV-GP transfected CHO-K1 cells

HEK-293 cells are commonly used as host for the heterologous expression of membrane proteins because they have high transfection efficiencies, faithfully process translation and glycosylation of proteins, and have optimal cell size, morphology and division rate. In addition to the HEK-293^EBOV-GP^ cells, we used EBOV-GP transfected CHO-K1 cells to assess the binding behavior of various recombinant fusion proteins. HEK-293 and CHO share very similar protein modification machineries in the endoplasmic reticulum (ER) and Golgi (103). More importantly, CHO cells are able to produce complex types of recombinant proteins with human-compatible glycosylation. CHO cells, however, do not express Gal a2,6 ST, a1,3/4 fucosyltransferase or b-1,4-N-acetylglucosaminyl-transferase III (GnT-III), which are enzymes expressed in human cells (104–106). Our results show, that proteins of the selectin-family (P-, L-, E-selectin) bind strongly to EBOV-GP transfected CHO-K1 cells. Siglec-7, NKp44 and NKp46 also bind significantly better to the transfected than to control cells, while PSGL-1 Siglec-3, Siglec-5, DC-SIGN and DC-SIGNR did not show significant binding compared to control cells (Fig 5). This is in agreement with the results of the ELISA - assays shown in Fig. 2.

**Fig. 5:**
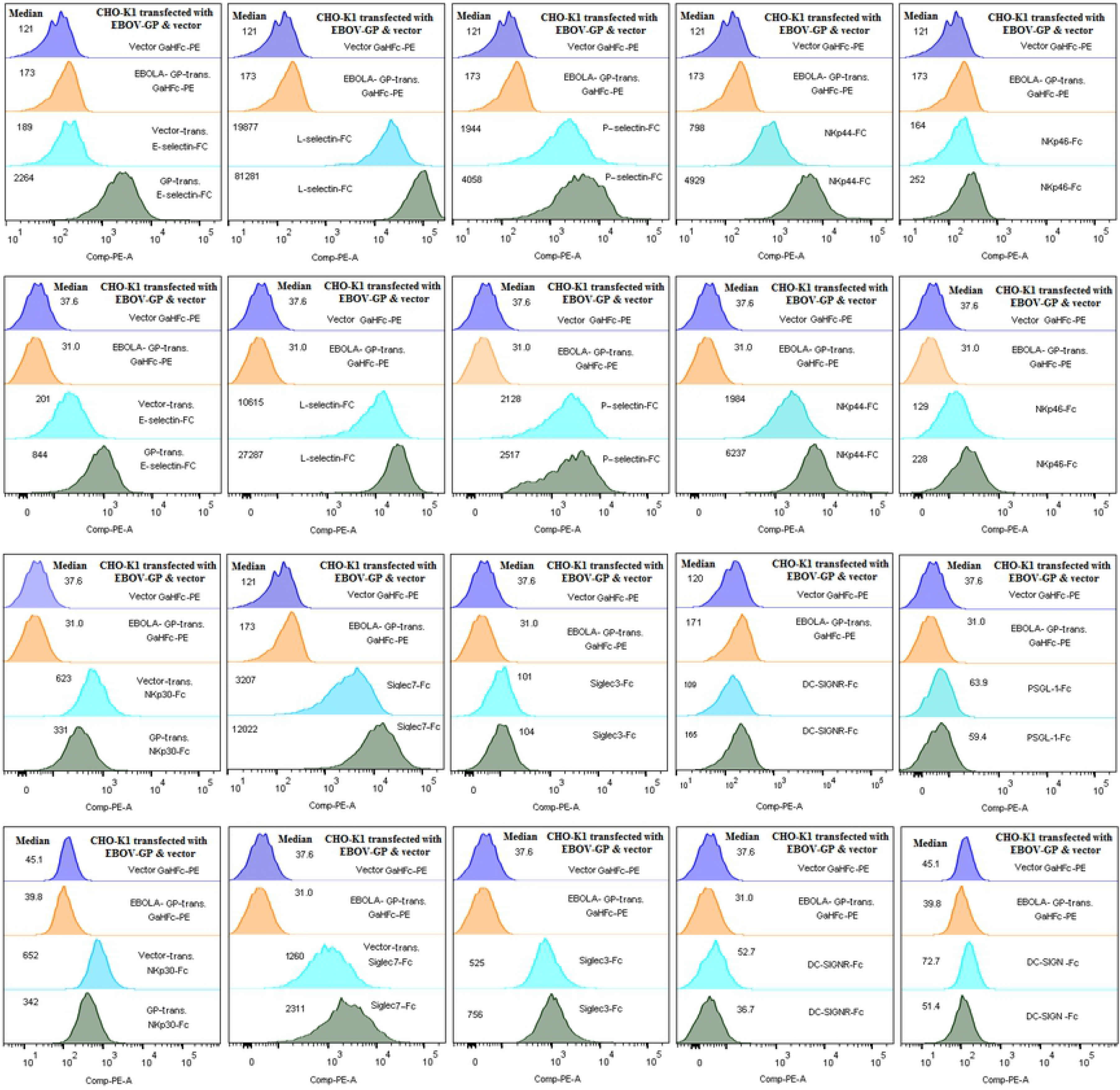
Binding of different recombinant fusion proteins to EBOV-GP transfected CHO-K1 cells compared to vector transfected controls. Shown are two independent experiments with similar results. L-selectin-Fc, E-selectin-Fc, P-selectin-Fc, Siglec-7-Fc, NKp44 and NKp46 have a higher affinity to the EBOV-GP transfected CHO-K1 cells. PSGL-1, Siglec-3, DC-SIGN and DC-SIGNR did not show affinity, and NKP30 showed even reduced binding to EBOV-GP transfected CHO-K1 cells, when compared to the respective controls.

### PSGL-1, CD24, CD44, DR3 and EBOLA-GP are receptors for selectins

The aforementioned selectins are able to interact specifically with CD24, CD44, DR3, PSGL-1, and LAMP1/2 (107–111). Flow cytometry analyses showed that binding of the CD44 monoclonal antibody to the HEK-293^EBOV-GP^ cells was increased compared to vector transfected cells, suggesting either upregulation of standard CD44 (CD44s) on the cell surface or steric hindrance of selectins binding CD44 in *cis* by competition with EBOV-GP. In HEK-293^EBOV-GP^ cells, EBOV-GP interacts on the cell surface in *cis* with selectins, resulting in more detectable CD24 and CD44 molecules.

CD24/ CD44 and EBOV-GP function as binding co-partners for P / E / L-selectins in *cis* and *trans*. *Cis* interaction of CD24 / CD44 and EBOV-GP expressed on the cell surface with selectins P / E-is not stable, hence there is an exchange between binding partners pending on their glycosylation. It is known that CD44 is hardly detectable on HEK-293 cells (112, 113). It should be considered that CD44 comprises a family originating from alternative splicing and undergoing posttranslational modifications with diverse functions, such as cell adhesion on different cell types (113). Proteoglycan CD44V6 (homing receptor Pgp-1, a variant of CD44) interacts with integrins and has been linked to poor prognosis in several tumors, where it facilitates metastasis and homing into distant tissues (114). We hypothesized that the ligands for selectins (CD44, CD24) on the cell surface of HEK-293^EBOV-GP^ cells probably are more accessible than on the vector transfected HEK-293 cells. Indeed, our results demonstrate an increased binding of the anti-CD44 and anti-CD24 antibodies to EBOV-GP transfected cells compared to vector transfected HEK-293 control cells (Fig. 6a).

**Fig. 6:**
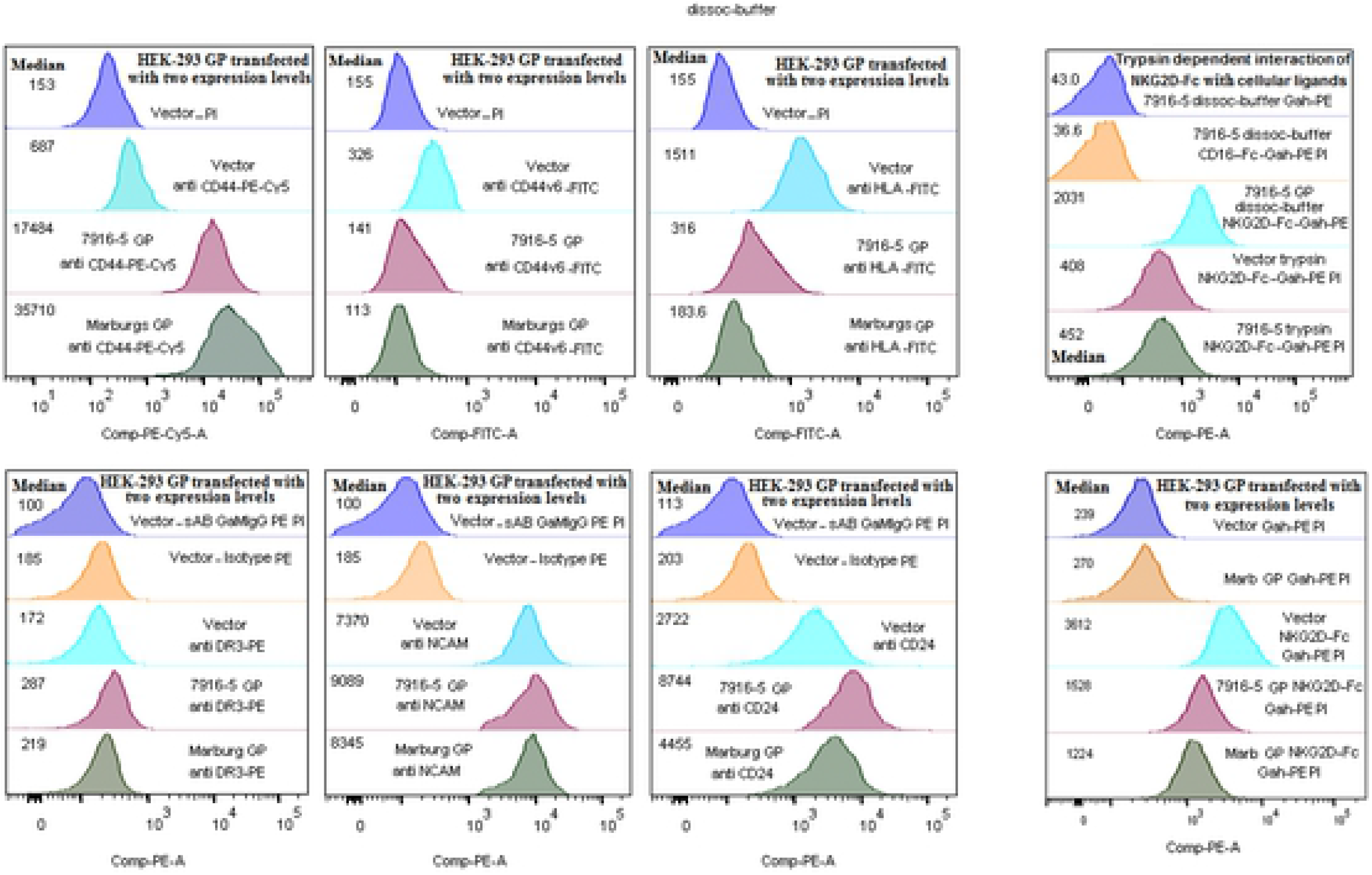
Detection of selectin receptors with antibodies and fusion proteins on HEK-293^EBOV-GP^ cells. (6a) Flow cytometry analysis showing the binding of different monoclonal antibodies against CD44v6, CD44s, DR3, CD24, HLAabc, and NCAM (CD56) to HEK-293^EBOV-GP^ cells compared to vector transfected controls. (6b), Binding of NKG2D-Fc to its ligands MICA/B (MHC I chain related molecule) is reduced in HEK-293 transfected cells either with the Marburg-GP or the 7916-5 GP, while binding of CD24 and CD44s is significantly increased. The expression of HLA is reduced. The binding of mAb against CD44v6, DR3 is not significantly changed.

When we incubated EBOV-GP transfected HEK 293 cells with CD24-Fc / CD44-Fc chimeric soluble receptors, which bind P / E / L-selectins, or with P / E / L-selectin-Fc chimeric soluble receptors, which bind CD24 and CD44, the ligand-receptor interactions in *cis* were disrupted (Fig. 6a). These biophysical interactions may also explain the increasing availability of CD24 / CD44 to bind anti-CD44s, anti CD44v6 and anti-HLA^abc^ on HEK-293 transfectants. The analysis showed that binding of NKG2D-FC to its cellular ligands MICA/B (MHC I chain related molecules) is reduced for both types of GP-transfected HEK-293 cells (see Fig. 6b), whereas CD24, CD44s and CD44v6 bindings are significantly increased. Furthermore, we observed that the binding of mAb versus DR3 was slightly increased, probably to support the apoptosis induction on effectors cells, while mAb against CD56 was not significantly changed. Our results with EBOV-GP transfected cells showed a reduction in MHC-I (Fig. 6a). In addition, we observed a lower binding of NKG2D-Fc chimeric soluble receptors to EBOV-GP transfected HEK 293 cells (Fig.6b), which indicates a reduction of its ligands MIC-A/B, as it has been shown before (115). Altogether, these data suggest that a by EBOV-GP reduced stimulation of NKG2D, together with the recruitment of Siglec-ligands on the surface of infected cells, may lead to a suppressive state of effector cells (Fig. 6b, bottom).

### Binding of the chimeric soluble receptors to their ligands depends on sialic acid and heparan sulfate

P- and L-selectin recognize clustered O-sialoglycan-sulfated epitopes on glycans, for example on the proteoglycan CD44. L-selectin, like other selectins, recognizes sialylated Lewis^x^ and sialylated Lewis^a^ (sLea; Neu5Acα2–3Galβ1–3(Fucα1–4)GlcNAc) (87, 116). All three selectins recognize sulfated and sialylated derivatives (116). In addition, L-selectin binds to O-glycosylated sialomucins (117). Many enveloped viruses, for instance influenza virus and Newcastle disease virus, bind to sialic acid residues located on the surface of target cells (72). For viral cell entry, a structural viral protein termed neuraminidase (F0) cleaves the glyosidic bond to sialic acid residues and thus unleashes the virus for fusion with the target cell membrane (59, 69, 71–73, 118). In order to analyze the involvement of sialic acid and heparan sulfate in the binding of the fusion proteins to EBOV-GP transfected CHO-K1 cells, we treated the GP and mock transfected cells with heparanases I & III, and sialidase. Following treatment, we analyzed the impact of these enzymes on the binding of fusion proteins to EBOV-GP and mock transfected CHO-K1 cells (Fig. 7). Treatment with heparanase resulted in a decreased binding of selectins to EBOV-GP transfected CHO-K1 cells. Sialidase treatment, on the other hand, showed a significantly higher binding of selectins to EBOV-GP.

**Fig. 7:**
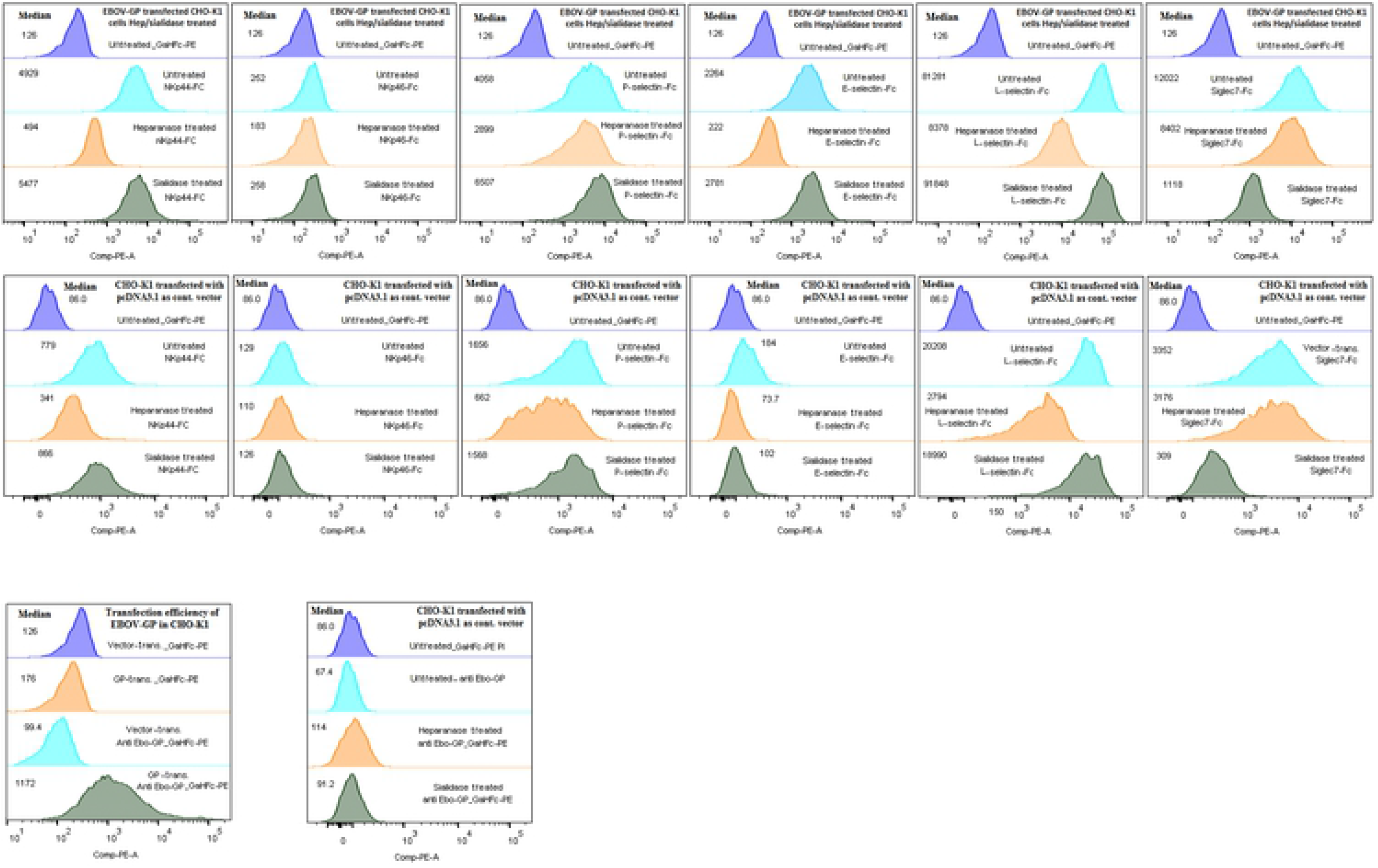
Treatment of EBOV-GP transfected CHOK1 cells with heparanase I & III and Sialidase. (7a) The binding of the fusion proteins to EBOV-GP transfected CHO-K1 cells changed significantly after removal of sialic acid and heparan sulfate. Treatment with heparanase resulted in decreased binding of selectins to EBOV-GP transfected cells. Sialidase treatment on the other hand caused a significantly increased binding of selectins to EBOV-GP. Binding of siglec7 was not affected by heparanase, yet strongly sensitive to sialidase I/III treatment, which reduced its binding significantly. This can be explained by the fact that sialic acid is the main ligand of siglecs. NKp44 and NKp30 have a reduced binding under heparanase treatment, as it is known that heparan sulfate is the main ligand for NCRs. (7b) Control of the transfection efficiency with EBOV-GP and the control transfected with empty vector.

The analysis of HEK-293^EBOV-GP^ cells showed inconsistent binding of the fusion proteins E-selectin-Fc and P-selectin-Fc, which could be explained by changes in the degree of sialylation and heparan sulfation of the cell surface proteins (fig 7). In agreement with this, we observed that different culture conditions of HEK-293^EBOV-GP^ cells changed the constellation and ratios of surface adhesion molecules and other glycoproteins, especially of selectin ligands (unpublished results and Fig. 4). CHO-K1 cells transfected with EBOV-GP, which were incubated with fusion proteins (NKp44-Fc, NKp46-Fc, and L / E / P-selectins-Fc) showed reduced binding following heparanase treatment, as was expected since it is known that heparan sulfate is the main ligand for NCRs (57, 58). Binding of Siglec7 was not affected by heparanase, yet strongly sensitive to sialidase I/III treatment, which reduced its binding significantly, as can be explained by the fact that sialic acid is the main ligand of Siglecs. These observations were consistent with the results obtained in the ELISA-assay (compare Fig 2 with Figs. 5 and 7).

### Binding of fusion proteins to lentiviral particles displaying EBOV-GP

The GP glycoprotein serves the Ebola virus to infect host cells by binding to various docking proteins. After transduction, viruses can incorporate or co-package many host cell-derived non-viral surface proteins into their newly formed envelope (119–121). Lately, this was demonstrated for infectious clones of HIV-1 propagated in HEK-293T cells (122).

Also, it was reported that TIM-1 interacts with GP Ebola virus glycoprotein (EBOV-GP) and phosphatidylserine (PS) on the viral envelope (dual binding) (88–91). However, in our study we could not prove binding of TIM-1 to the HEK-293^EBOV-GP^ cells or purified EBOV-GP protein. Nevertheless, we could show binding of TIM-1 to lentiviral particles displaying EBOV-GP on their envelope (lenti-EBOV-GP).

The recombinant lentiviral particles were produced in HEK-293 cells stably transfected either with a plasmid expressing EBOV-GP (pcDNA6/V5-HisB-EBOV-GP) or just vector (pcDNA6/V5-HisB). After concentrating the lentiviral particles, they were used for coating ELISA-plates. The efficiency of coating was determined using anti-GP-hIgG-Fc and goat anti-hIgG-POX. The coated plates were incubated with the most relevant fusion proteins. The results are shown in Fig. 8. We found significant binding to lenti-EBOV-GP by NKp44, NKp46, CD62e/l/p (E-Selectin, L-Selectin, P-Selectin), inhibitory receptor siglecs (siglecs-5, siglecs-7), DC-SIGN, and DC-SIGNR. From these, Siglec-3 and -5 showed weaker binding. In contrast, NKp30, PSGL-1, CD44, CD24, syndecan-1^1^, and TIM-1 did not bind to the Lenti-EBOV-GP. The observed binding of DC-SIGN, DC-SIGNR and selectins to the lentiviral particles without EBOV-GP is in agreement with previous reports (75, 77, 81). The binding of TIM-1 to lentiviral particles irrespective of their content of EBOV-GP is due to the fact that TIM1 binds to cohesive phosphatidylserine, which originated from the cell surface of host HEK-293 cells transfected with vector or EBOV-GP, and were taken up by the virus envelope (120) (Fig. 8). Likewise, the binding of CD44 and PSGL1 fusion proteins to vector envelopes, independent of any GP presence, can be explained by their binding to cellular ligands from the host plasma membrane, which were taken up by the lentiviral envelopes. The above described interaction plays a role in the recognition of virus infected cells by immune effector cells, e.g. NK cells. Interestingly, besides these specific interaction partners, a second mechanism of interaction may occur. This is the interaction of other cellular targets, e.g. membrane-associated heparan sulfate proteoglycans, which are recognized by NKp30, NKp44 and NKp46 (123) from NK-cells.

**Fig. 8:**
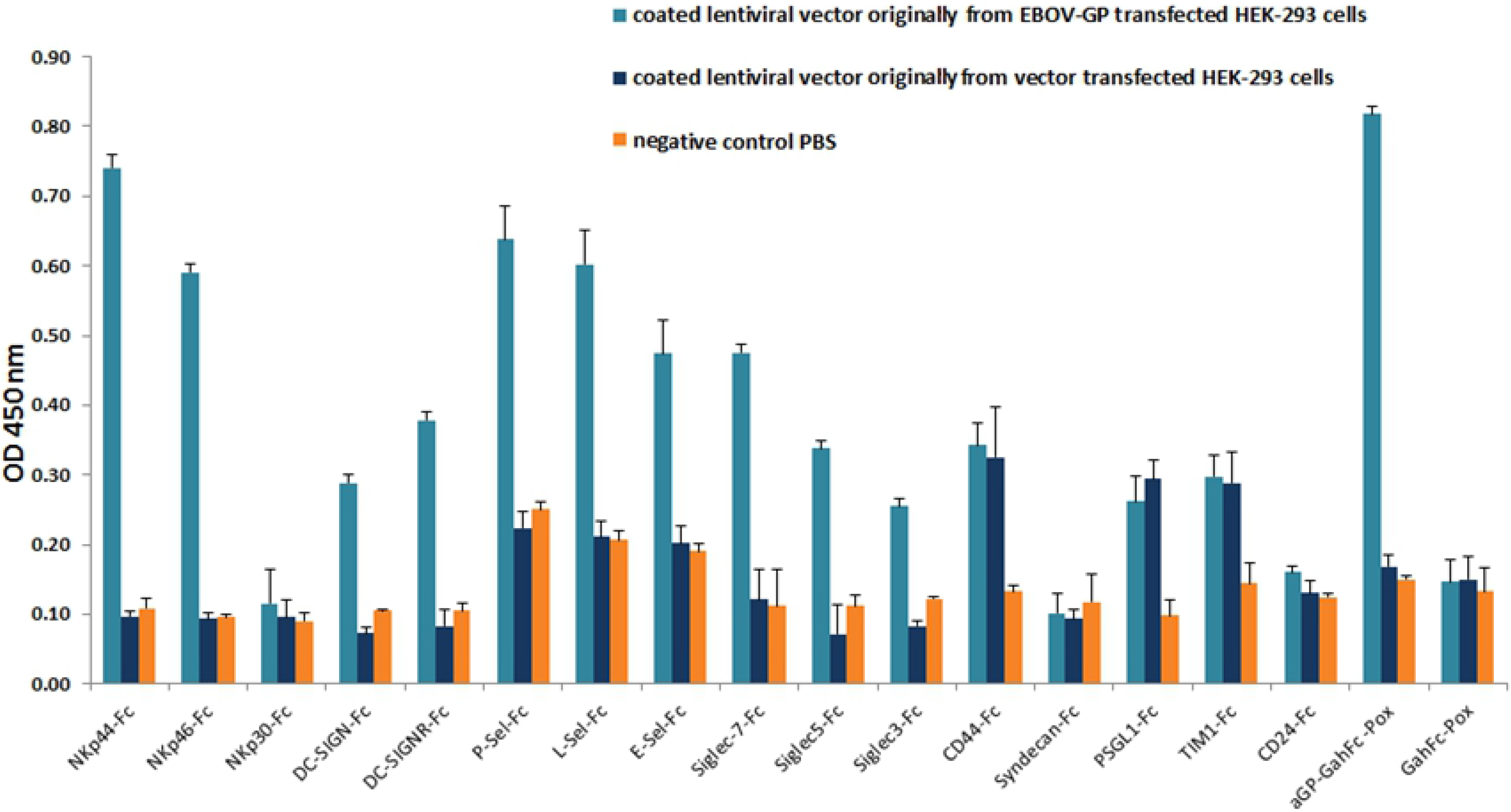
Binding of fusion proteins to lentiviral particles displaying EBOV-GP. The lentiviral particles were used for coating ELISA-plates. The coating efficiency was determined by monoclonal antibody KZ52 and secondary antibody GahIgG1-POX versus lentiviral vector with or without GP glycoprotein of Ebola virus and compared to the corresponding lentiviral vector control. The results show significant binding of NKP44-Fc, NKp46-Fc, E-selectin-Fc, L-selectin-Fc, E-selectins-Fc, and Siglec7-Fc. There was reduced binding of Siglec3-Fc and DC-SIGNR (CD299)-Fc to the GP containing vector lentiviral particle. There was equally increased binding of CD44-Fc PSGL-1-Fc and Tim-1 to the lentiviral vector with or without GP.

### HPV-L1 binds NCRs, Homing Selectins and inhibitory Siglecs

Human papillomavirus infection requires cell surface heparan sulfate. The first interaction of HPV virions with the epithelial cell surface is with heparan sulfate proteoglycans (HSPGs) as it was shown for HPV16 and HPV33 pseudovirions in COS-7 cells (124). Furthermore, expression of the integral membrane proteins syndecan-1 and syndecan-4 in K562 cells, which do not have common HSPGs, was shown to enhance the attachment of HPV16 L1 virus-like particles (VLPs) and HPV11 virions in good correlation with the expression levels of HSPGs. In contrast, GPI-anchored glypicans are not primary receptors for binding of HPV VLPs to keratinocytes (125). In order to further asses the binding of the fusion proteins to purified HPV-L1 virus-like particles coated on ELISA plates, we incubated them with the different fusion proteins. Our results suggest that especially NKp44, P-selectin, and L-selectin bind strongly to HPV-L1, while NKp46, Siglec-5 and Siglec-7 showed lower binding (Fig. 9). Out of the group of selectins, L-selectin showed the highest binding affinity to HPV-L1. Although we had HPV-L1-transfected HEK-293 cells, we could not use them to test binding of our fusion proteins, as HPV-L1 does not have a transmembrane domain and therefore is secreted.

**Fig. 9:**
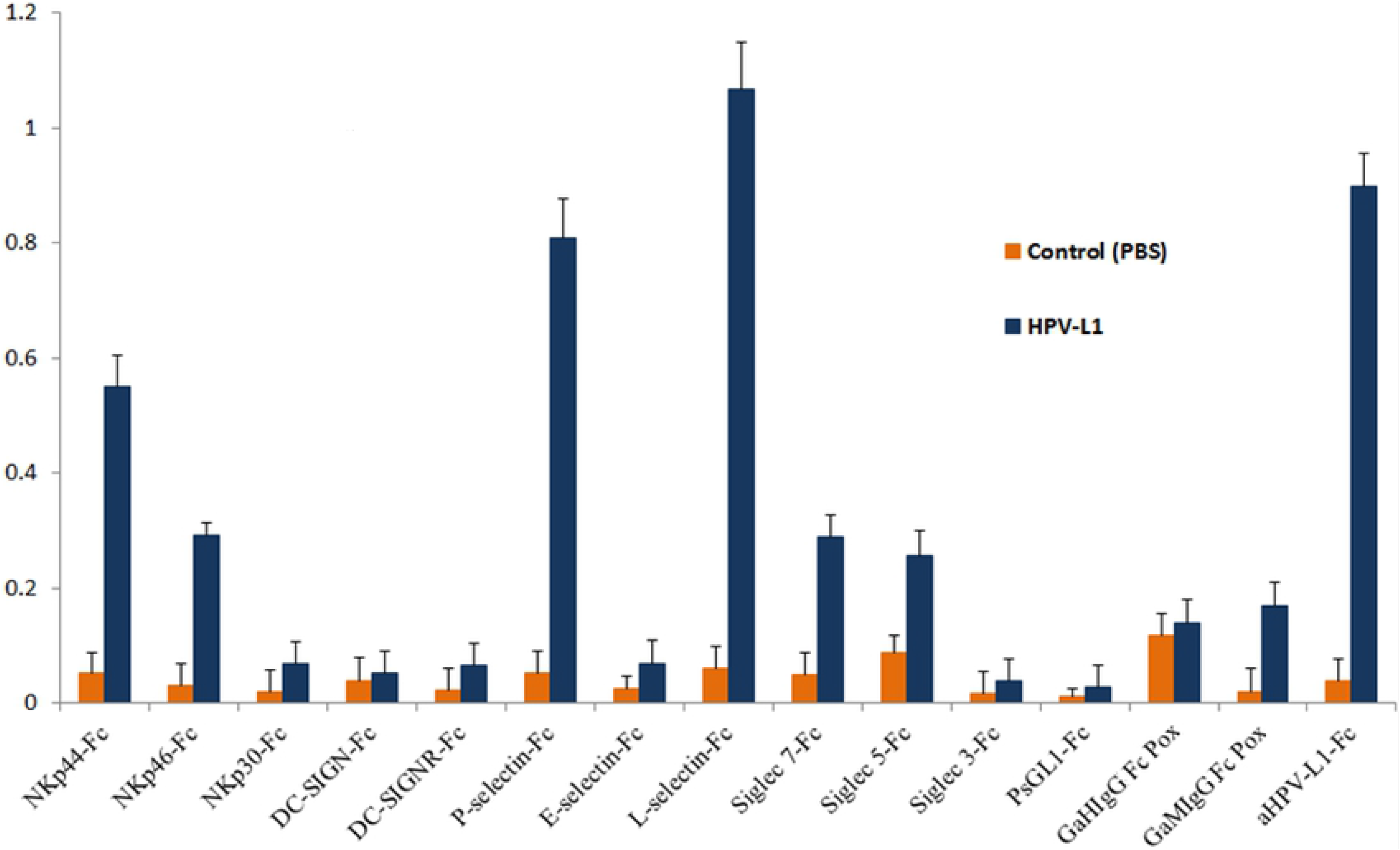
Binding of chimeric soluble proteins to purified HPV-L1 particles. HPV-L1 coated ELISA-plates were used to analyze the binding of the indicated chimeric soluble proteins. Goat-anti-hIgG-POX and Goat-anti-mIgG-POX served as secondary antibodies. The fusion proteins with PBS-coated plates served as negative controls. For positive controls, we used anti-HPV-L1 and secondary anti-mIgG-POX for detecting purified HPV-L1 on coated ELISA-plates. NKp44, NKp46, P-selectin, L-selectin, and inhibitory receptor Siglecs (Siglec-5, Siglec-7) bind significantly to HPV-L1. NKp30, PSGL-1, E-selectin, Siglec-3, DC-SIGN (CD209) and its receptor DC-SIGNR (CD299) did not show specific binding. This experiment was repeated four times under different blocking conditions.

### EBOV-GP protects HEK-293 cells from lysis by polyclonal NK cells

We have observed that GP expression by target cells reduced lysis caused by primary NK cells, although staining of MHC-I by W6/32 mAb was significantly reduced (Fig.6a). Thus, this unexpected reduction could result from masking HLA-I by EBOV-GP. In this case, however, increased NK-cell activity could be anticipated. However, we reasoned that other mechanisms are responsible for the reduced killing of target cells by NK-cells. Therefore, we assessed the susceptibility of HEK-293^EBOV-GP^ cells to lysis by primary NK cells obtained from different donors in comparison to respective mock-transfected HEK-293 cells. As shown in Fig 10b, HEK-293^EBOV-GP^ cells were significantly less susceptible to lysis by activated polyclonal NK cells compared to mock-transfected cells, suggesting that the pleiotropic effects of EBOV-GP on the cell surface have an overall suppressive result on NK cells, most likely by reduction of NKG2D ligands and by interacting with Siglec7 and TIM receptors. Furthermore, monoclonal antibodies directed toward NCRs reduce the activity of NK cells (Fig. 10a). Altogether, these results suggest a crucial role of EBOV-GP in mediating immune escape of transduced/infected cells.

**Fig. 10:**
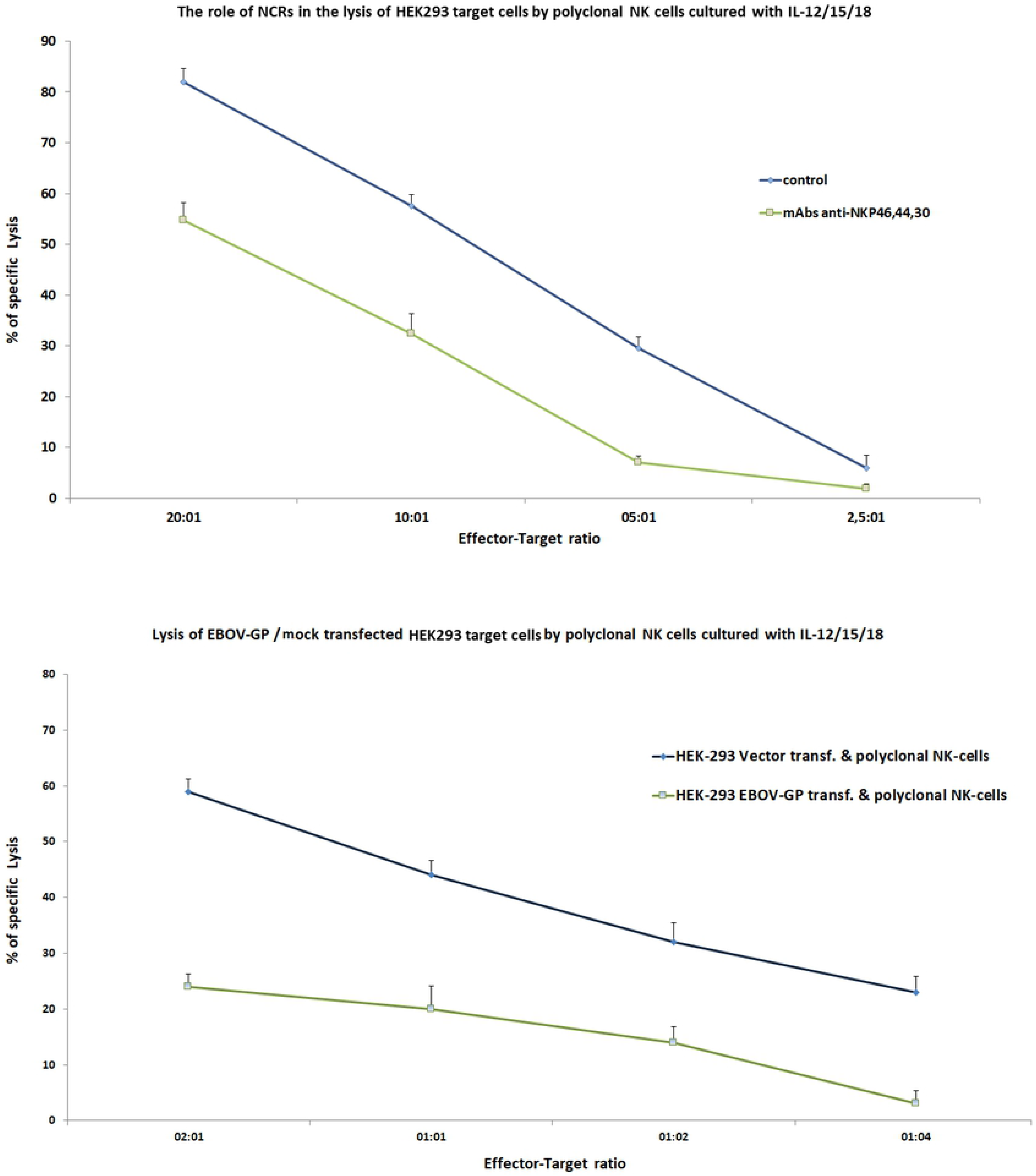
Lysis-reduction of HEK-293^EBOV-GP^ target cells by polyclonal NK-cells. (10a) NK activation receptors NKp44, NKp46, inhibitory Siglecs and L-selectin are able to bind strongly to HEK-293^EBOV-GP^ cells, compared to mock transfected HEK-293 cells. The results of a chromium release assay indicate that HEK-293 target cells show decreased susceptibility to lysis by polyclonal NK-cells. In this assay, specific anti-NCRs antibodies blocked the interaction between NCRs and their ligands on HEK-293 cells. (10b) Results of a chromium release assay using HEK-293 target cells transfected with EBOV-GP show decreased susceptibility to lysis by polyclonal NK-cells in comparison to non-transfected HEK-293 cells.

**Fig. 11:**
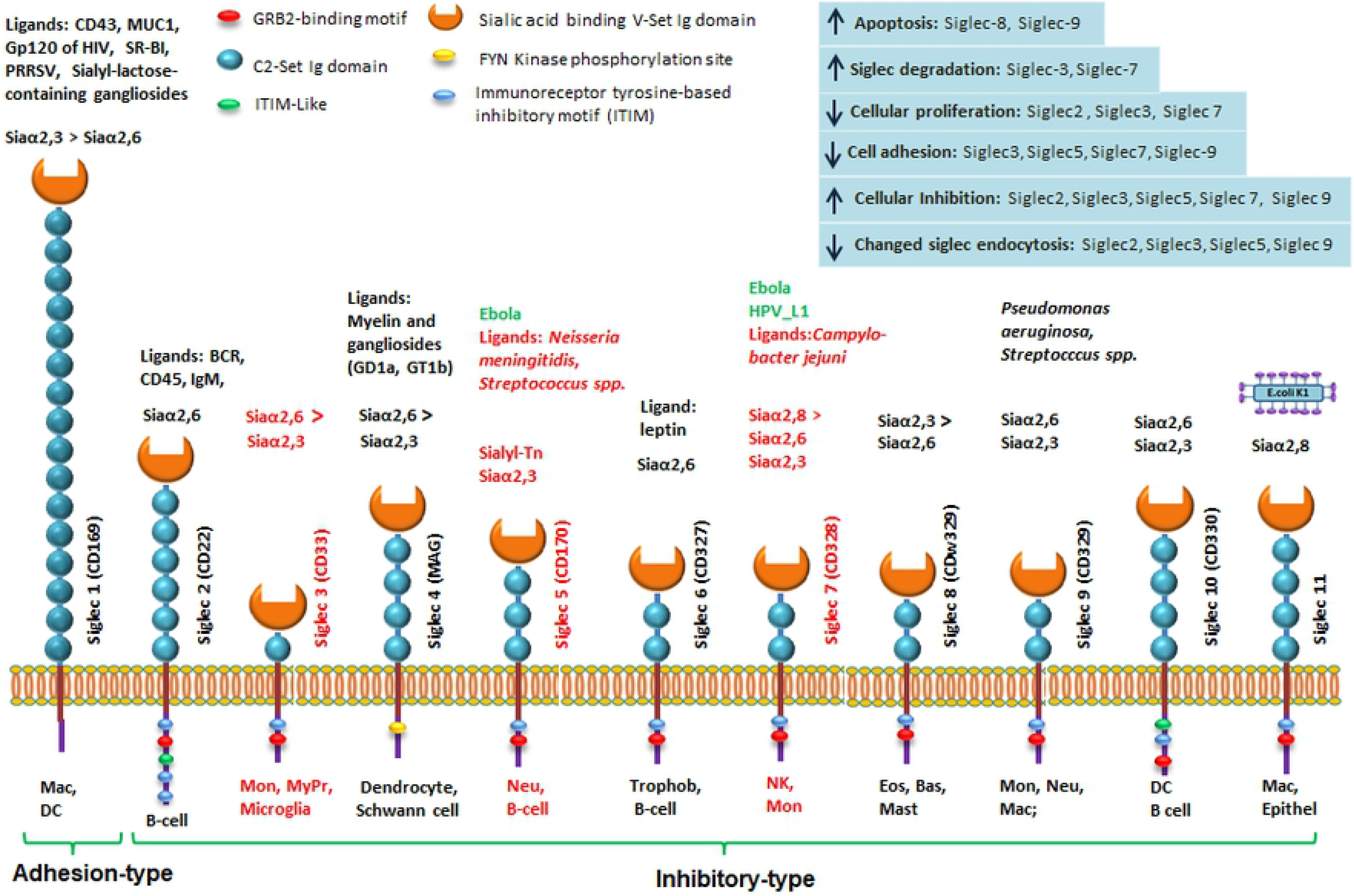
Specific relations of Siglecs with their sialylated ligands and resulting functions. The important Siglecs are shown with their intracellular-, transmembrane-, and extracellular domains (bottom part). The Siglecs’ roles depend on their specific binding domain (V-Set Ig domain) as well as on intracellular inhibitory motives (GRB2, ITIM-like, ITIM, FYN kinase phosphorylation site; see legend). Cells, which express the various Siglecs are given below the respective symbols (Mac: macrophage, DC: dendritic cell, Mon: monocyte, MyPr: myeloid precursor, Neu: neutrophilic cell, B: B-cell, Trophob: trophoblast cell, NK: natural killer cell, Eos: eosinophilic cell, Bas: basophilic cell, Mast: mast cell, Epithel: epithelial cell, Osteoclast: osteoclast cell). Examples of ligands binding to the Siglecs are indicated above their symbols. Gp120 of HIV: glycoprotein 120 from HIV capsid, SR-BI: Scavenger receptor class B type 1, PRRSV: porcine reproductive and respiratory syndrome virus, MUC1: Mucin 1, CD43: semaphorin, Sialyl-lactose containing gangliosides, Siaα2,3 > Siaα2,6: Preferred sialic acid linkage type of respective binding partners. Further ligands of other siglecs: BCR: B-cell receptor, CD45, IgM, GD1: ganglioside 1, GT1b: ganglioside, carbohydrates from selected gram-negative bacteria (e.g. Neisseria meningitidis, Campylobacter jejuni, Pseudomonas aeruginosa, E.coli K1, etc.) and some gram-positive bacteria (e.g. Streptococcus spp.) that express sialic acid. In addition, the preferential sialic acid linkages are indicated.

## Discussion

NK cells are effector cells of the early innate immune response that play a critical role in the lysis of virus infected and tumor cells without requiring prior antigen stimulation (126, 127). The functions of NK cells are regulated through a balance of activating and inhibitory signals, which are transmitted through particular receptor binding cytokines or ligand structures on interacting target cells and pathogens (52, 128–131). Activating signals of NK cells lead to either exocytosis of cytotoxic granules that lyse the target cells (132–135) or release of extracellular ligands, including Fas ligand (FasL) or tumor necrosis factor-related apoptosis-inducing ligand (TRAIL) (136). Stress, which is caused by altered metabolism, virus infection or transformation, can enhance expression of activating ligands and decrease expression of inhibitory ligands (MHC-I), which transfers affected cells to targets of innate immune effector cells. In murine or human T cells, inhibitory KIR binding to MHC-I of target-cells enhances the survival of CD8+ T cells (137). The activating ligands are independent of antigen presentation and can mobilize CD8^+^ T, NKT, and NK effector-cells. Ebolavirus uses evasive mechanisms to directly disrupt antiviral effector cells of the adaptive and innate immune system (NKT, CD8^+^ T and NK cells).

We decided to study the mechanism of this process and the limitations of the immune effector cells to eradicate Ebolavirus. It is our notion, that the elimination of virus infected target cells by NK-cells is based on the sum of their ligand-binding activities, which is determined by the integration of inhibitory (e.g. KIRs) and activating receptor signals. We and others have previously shown the interaction of NCRs with hemagglutinin-neuraminidase of Newcastle disease virus (NDV), poxviral hemagglutinin and influenza viruses (59, 69, 72). We hypothesized that activating NCRs play a key role in the recognition of structural glycoproteins of virtually any virus by NK effector cells and thus facilitate the elimination of pathogens. Therefore, we have investigated the binding of purified EBOV-GP and HPV L1 to a series of soluble ectodomains of NCRs, selectins, Siglecs and other homing receptor proteins fused to the Fc portion of human IgG1. Recently, it was reported that blocking Nkp30 by a specific antibody reduced lysis of Ebolavirus infected dendritic cells by NK effector cells (30). So far, it is unknown, to which degree the cell surface of virus-infected cells is changed by the presence of EBOV-GP and then can sterically hinder the interaction of host cell adhesion proteins with activating or inhibiting effector-cell receptors. Nevertheless, it is interesting that EBOV-GP expressing cells can directly switch off the function of NK and CD8 T cells (138–140). The mechanism by which EBOV-GP suppresses the activity of NK cells may simultaneously involve (i) the binding of different inhibitory Siglecs, (ii) a reduction in ligands of KIR-receptors (MHC-I), some of which have activating roles, such as those KIRs with short cytoplasmic domain associated with activating adaptors (e.g. 3DS1 and 2DS1-5) and (iii) a reduction of NKG2D ligands (MICA/B). Specifically, highly glycosylated viral EBOV-GP protein leads to reduced expression or shedding of MHC-I in host cells, or acts in *cis* sterically so that interaction or polarization between virus-infected target and effector cells is blocked (22) (Fig 6a/b). Besides KIR family genes, NK cells encode a variety of other activating and inhibitory receptors. In addition, there is inter-individual genetic diversity that causes different levels and types of inhibitory (KIR) and activating receptors (NKG2D, NCRs) (59). Here, we observed a lower binding of NKG2D-Fc chimeric soluble receptors to EBOV-GP transfected HEK 293 cells corresponding to reduced expression of MICA/B (Fig. 6b, bottom), as has been reported before (115). Interestingly, we reported previously for Vaccinia virus (VV)-infected HeLa cells that the cumulative expression of NKG2D ligands (MICA, MICB, ULBP1, ULBP2, ULBP3, and ULBP4) was identical to that in non-infected cells, as detected with NKG2D-Fc (59, 141, 142). However, a detailed analysis of the NKG2D ligands revealed a differential modulation of single ligands (59). In contrast, infection of HeLa cells with the mouse poxvirus ECTV resulted in clearly reduced levels of the cumulative expression of NKG2D ligands, as detected by human NKG2D-Fc (59). These data suggest that a by EBOV-GP reduced stimulation of NKG2D, together with the recruitment of Siglec-ligands on the surface of infected cells, may lead to a suppressive state of effector cells.

Furthermore, our data suggest that EBOV-GP binds strongly or at least moderately to the chimeric soluble proteins L-and P-selectin, Siglec-7 and -5, NKp44, NKp46 and, to a lesser extent, to Siglec-3 (Figs. 2, 4, 5, 7, 8). DC-SIGN, DC-SIGNR, and TIM-1 are known cellular receptors for EBOV (88, 90, 143–145). Additionally, we investigated binding of DC-SIGN-Fc, DC-SIGNR-Fc and TIM-1-Fc to HEK-293^EBOV-GP^ cells and to purified EBOV-GP coated on ELISA plates. Interestingly, we were able to demonstrate the binding of DC-SIGN and DC-SIGNR to purified EBOF-GP on ELISA plates, but not to HEK-293^EBOV-GP^ cells. A possible explanation is that EBOV-GP from the surface of transfected cells is fully occupied by interaction with its target proteins in *cis*, which causes that the binding sites for DC-SIGN-Fc and DC-SIGNR-Fc are not free for interacting in *trans* (Figs 2, 4, 5 and 8). In contrast, GP on the viral envelope remains free for interaction with both proteins. Like the HIV-1 gp120, the highly glycosylated EBOV-GP utilizes the C-type lectin receptor DC-SIGN (CD209) to infect dendritic cells, which are a major reservoir of EBOV (143–145). Soluble GP is post-translationally modified by N-glycosylation, but importantly also by C-mannosylation (9). DC-SIGNR specifically interacts with high-mannose N-linked carbohydrates on viral pathogens.

Also, we could not prove binding of TIM-1-Fc to HEK-293^EBOV-GP^ cells or to purified EBOV-GP, yet we showed binding of TIM-1-Fc to lentiviral particles displaying EBOV-GP on their envelope (lenti-EBOV-GP), as well as to mock lentiviral particles. The binding of TIM-1-Fc to lentiviral particles irrespective of their content of EBOV-GP is due to the fact that TIM-1 binds to cohesive phosphatidylserine, which was incorporated by the virus envelope. It originates from the inner plasma membrane of mock or EBOV-GP-transfected HEK-293 cells (120) (Fig. 8). TIM-3, TIM-4, and TIM-1 together inhibit HIV and Ebola virus release from infected cells (146). TIM-1 also serves as a pattern recognition receptor on invariant natural killer cells (iNKT), which mediate cell activation by TIMs binding to phosphatidylserine on the surface of cells undergoing apoptosis (147). It is very likely that TIM-3, an inhibitory checkpoint receptor on membrane effector cells, binds also to phosphatidylserine of the EBOV envelope (92, 93). EBOV can induce dysfunction and even apoptosis in effector cells by binding to Siglecs-7,-5,-3,-8,-9, TIM-3 (91, 148–151) and probably to other inhibitory receptors and thus cause immune escape (91, 152, 153). In line with this, a deficiency of immune effector cells has been observed during EBOV infection, whereby virus replication continues to be uncontrolled and renders immunotherapy of the disease difficult (154, 155). The strength of NK cell cytotoxicity is controlled by an interplay of inhibitory (KIR, Siglecs, checkpoint proteins) and activating receptors (NCRs, NKG2D), as well as various costimulatory molecules (CD28, CD2) (156, 157). A critical threshold of activation must be achieved for NK cells to mount a productive response (158, 159). Viral infection can lead to immune suppression, either by downregulation of the cytotoxic function or by triggering apoptosis, leading to depletion of effector cells. Surprisingly, studies on EBOV-GP infected animals have shown a steady decrease in NK cell activity and the number of NK cells (140). It is unclear, to which extent adhesion molecules, which play important roles in interacting with the activating or inhibiting NK cell receptors, are affected by EBOV-GP in their relative expression, or through steric hindrance. More in-depth knowledge about the mechanisms by which EBOV-GP expressing cells are able to directly switch off NK cell functions will certainly be helpful to develop an efficient therapy for EBOV infections (140).

Siglecs often function as sensor for sialylated glycoproteins. Through their intracellular ITIM, they induce strong inhibitory signaling upon binding to different linkages of sialic acid (160). Interestingly, this mechanism is used by tumor cells and pathogens to escape the immune system, by adding sialic acid residues to their glycan structures, thus highlighting that the sialic acid-Siglec interaction is key to the immune function against pathogens and cancer (161, 162). NK and other effector cells express various Siglecs, e.g. Siglec-3, 7, 8, and 9 (163, 164). Their presence stabilizes the conformation of membrane glycoproteins in *cis* interactions with endogenous sialo-conjugates at the cell surface of NK cells (164). Accordingly, we have observed binding of Siglec-7 and -5 to EBOV-GP in ELISA as well as to CHO-K1 EBOV-GP transfected cells and lentiviral particles. Siglec-7 binding to EBOBV-GP transfected CHO-K1 is dependent on 2,6 sialic acid linkage, since treatment with sialidase reverts the binding (Fig. 7). We have not tested Siglec-8 and -9, but strongly believe that Siglecs-8 and -9 also bind to GP because of their similar affinity for 2,6 and 2,3 sialic acid linkages. In addition, they are MHC class I-independent inhibitory receptors on immune effector cells, which–when stimulated-prevent the activation of these cells. However, the expression of the corresponding ligands on target cells results in inhibition of NK cell-mediated cytotoxicity or apoptosis by interacting with Siglec-7 and -9 (164). The outermost position of sialic acids during the glycosylation process implies the capping of a variety of glycosylation structures (165). Indeed, sialic acid is increased on the surface of infected and tumor cells, which enables resistance against innate and adaptive immune responses and interference with the complement system (30).

Heparan sulfate binds many microorganisms and interacts with many viral envelope components e.g. HIV-1 (166), Hepatitis viruses (167, 168), Flavioviruses (169, 170), Vaccinia virus (171), human papillomavirus (HPV) (172), human herpes virus (173), HSV-1 (174), Ebola virus (175), Corona virus and so on. Heparan sulfate proteoglycans (HSPG) are composed of a core protein covalently linked to heavily sulfated and negatively charged glycosaminoglycan (GAG) chains and facilitate virus entry and attachment.

All selectins (E-, L- and P-Selectin) bind specific fucosyl-sialyl-Lewis group carbohydrates of surface glycoproteins such as CD44v6, CD24, PSGL-1, and CD43 (87). CD44 is a receptor for hyaluronic acid, which is commonly expressed in hematopoietic cells, fibroblasts, and many tumor cells, which contributes to the adhesion of these cells to the extracellular matrix. Hyaluronic acid degradation fragments can disrupt the interaction between different CD44 variants and hyaluronic acid (176). Such disruption affects cell-cell and cell-matrix interactions, including cell traffic, immune cell homing and inflammation, cell aggregation, and release of chemokines or growth factors (177). In contrast to normal cells, tumor cells with upregulated CD44v6 are not able to bind hyaluronic acid. Unlike all other isoforms, CD44v6 is decorated by a fucosyl-sialyl-Lewis^a^ group, which is transferred by fucosyltransferase 3 and this specific change blocks hyaluronic acid binding to CD44v6 (178, 179). We have shown that L-, P- and E-selectins bind to purified EBOV-GP coated on ELISA plates and to EBOV-GP transfected cells (HEK-293 and CHO-K1). This is confirmed by recently published data that the viral particle gp120 of HIV-1 binds L-selectin (CD62L) (84–87). Similarly, human P-selectin glycoprotein ligand-1 is a functional receptor for enterovirus 71 (180).

We conclude that selectins play an important role in virus release from infected cells, as it has been shown for HIV, which binds L-selectin and CD34 to co-localize with ADAM17 (84–87). Glycans designated sialyl 6-sulfo Lewis^x^ carbohydrate are potential ligands for selectins [15]. The interaction of the C-type CRD in the extracellular portion of each selectin with its receptor involves the direct binding of a fucose residue in the sialyl Lewis^x^ tetrasaccharide to the conserved calcium ion characteristic of the C-type CRDs [11,12]

The binding of NCRs to EBOV-GP as well as other viral envelope proteins, such as NDV, POX, HCMV, and Influenza, (59, 68–73, 181, 182) led us to examine their binding also to HPV-L1. Therefore, we tested the binding of the chimeric soluble proteins to HPV L1 virus-like particles. NKp44, P- and L-selectins showed strong binding to HPV-L1, while NKp46, Siglec-5 and -7 showed low binding (Fig. 9). Out of the group of selectins, L-selectin was observed to have the highest binding affinity to HPV-L1. Based on our results it is conceivable, that NCRs will also bind to the spike of Corona virus, as was suggested recently^2^.

## Materials and Methods

### Cell lines

FACS was used to analyze binding of recombinant fusion proteins to EBOV-GP transfected cell lines including human embryonic kidney cells HEK-293 (ATCC CRL-3216), HeLa cervix carcinoma cells (ATCC^®^ CCL-2™), and CHO-K1 (Chinese-Hamster Ovary ATCC ^®^ CCL-61™)^3^. Cell lines were cultured in RPMI 1640 (Invitrogen, Karlsruhe, Germany) supplemented with 2 mM glutamine and 10% fetal calf serum (FCS). Human polyclonal NK cells were isolated by NK cell negative isolation kits (Invitrogen) from peripheral blood or from normal donor buffy coats obtained from the blood bank. Between 95 and 99% of NK cells were CD3 negative and CD56 positive. Cells were grown in Iscove’s modified Dulbecco’s medium (Invitrogen) with 10% human serum, penicillin-streptomycin, and 100 IU/ml IL-2 (NIH Cytokine Repository, Bethesda, MD).

### Transfections

From the Institute of Virology in Marburg, Germany, we kindly received the vector pCAGGS-EBOV-GP for transfecting HEK-293-cells, which contains an AG promoter (chicken b-actin / rabbit b-globin hybrid). Furthermore, we used a blasticidin-expression vector (pcDNA6/His A) for HEK-293 cells, while CHO- and HeLa cells were transfected with a neomycin (G418) containing vector (pCDNA 3.1). Both contain a CMV promotor. The cDNAs were subcloned into the expression vector pcDNA3.1 (Invitrogen), pCAGGS.cm5 (Invitrogen), and pcDNA6 to generate pcDNA3.1(+)/GP, pcDNA6(+)/GP and pCAGGS.cm5 (+)/GP respectively. HEK-293 cells, CHO-K1 and HeLa cells (2.5 × 10^5^ respectively) were cultured in six-well plates and transfected with 4 μg of pcDNA3.1(+)/GP, pcDNA6(+)/GP, pCAGGS(+)/GP or the mentioned empty plasmids as vector and 10 μl of lipofectamine 2000 (Invitrogen) according to the manufacturer’s instructions. Two days later, cells were selected with geneticin or blasticidin (1 mg/ml in Dulbecco’s phosphate buffered saline [D-PBS]; Sigma-Aldrich, Taufkirchen, Germany) and, after being sorted for high-level GP expression, maintained in 0.5 mg/ml geneticin. GP expression was detected via mouse monoclonal anti-ZEBOV GP antibody 3B11 (102) (kindly provided from Prof. Stephan Becker, Institute of Virology, Marburg), and human monoclonal anti-ZEBOV GP antibody KZ52 (101) (dilution 1:200). KZ52 was purchased from IBT Bioservices (Integrated Biotherapeutics Inc, North Potomac).

### Chromium release assay

EBOV-GP transfected HEK-293 cells, which were used as target cells (0.5 × 10^6^) in 100 μl of assay medium (Iscove’s modified Dulbecco’s medium with 10% FCS and 1% penicillin-streptomycin), were labeled with 100 μCi (3.7 MBq) of ^51^Cr (Hartmann & Braun, Frankfurt, Germany) for 1 h at 37°C. Cells were washed twice and resuspended in assay medium at 5 × 10^4^ cells/ml. Effector cells were re-suspended in assay medium and mixed at different effector-to-target cell ratios with 5,000 labeled target cells/well in a 96-well F-bottom plate. Maximum release was determined by the incubation of target cells in 1% Triton X-100 solution. Spontaneous ^51^Cr release was measured by incubating target cells in the absence of effector cells. All samples were prepared in triplicate. Plates were incubated for 4 h at 37°C. Supernatant was harvested, and ^51^Cr release was measured in a γ-counter. The percentage of cytotoxicity was calculated according to the following formula: [(chromium release for condition of interest – chromium release in spontaneous wells)/ (max chromium release – chromium release in spontaneous wells)] × 100. The ratio between the amounts of maximum and spontaneous release was at least 3 in all experiments. Representative examples from two to three similar experiments are shown.

### ELISA-assay

For the direct detection of EBOV-GP via ELISA plates, we obtained the human Ebola Zaire (H. sapiens-wt) glycoprotein from Advanced Biomart (San Gabriel, CA, USA) and the anti-Ebola surface glycoprotein [KZ52] from Absolute Antibody (Oxford, GB). The obtained EBOV-GP was expressed with a polyhistidine-tag at the C-terminus and consists of 629 amino acids, predicting a molecular mass of 69.3 kDa. This antibody detects purified EBOV-GP coated on ELISA-plates and HEK-293^EBOV-GP^cells by IF. MicroTest III enzyme-linked immunosorbent assay (ELISA) plates (BD Biosciences, Heidelberg, Germany) were coated overnight with EBOV-GP in 0.05 M NaHCO_3_-Na_2_CO_3_ buffer (pH 9.6). They were also blocked using 3% skim milk powder (Merck, Darmstadt, Germany) in PBS-0.05% Tween 20 (Sigma-Aldrich) (PBS-T), as well as Pierce™ Protein-Free (PBS) Blocking Buffer (Thermo Scientific), overnight at 4° C. The recombinant fusion receptors listed above were obtained from R&D (Germany) and analyzed regarding their binding ability to EBOV-GP coated ELISA-plates. All purified recombinant proteins (1 μg/100 μl in PBS-T with 1% bovine serum albumin) were added to EBOV-GP-coated wells at 2μg/well for 1 h at room temperature. After three times washing with PBS-T, peroxidase-conjugated goat anti-hIgG Fc or goat anti-mouse IgG Fc in PBS-T (1:2,000 with 1% bovine serum albumin) were added for 1 h at room temperature and then the plates were washed three times with PBS-T. Thereafter, a peroxidase substrate solution (o-phenylenediamine [Sigma-Aldrich] at 1 mg/ml in 0.1 M KH_2_PO_4_ buffer [pH 6.0]) was added for 20 min at room temperature in the dark. The substrate reaction was stopped with 50 μl of 4 N H_2_SO_4_, and results were read with a Titertek Multiscan plus MKII ELISA photometer (MP Biomedicals, Heidelberg, Germany) at OD450 nm and 570 nm for reference.

### Flow cytometry

For cell surface immunofluorescence staining, 0.5 × 10^6^ to 1 × 10^6^ cells were washed once in ice-cold fluorescence-activated cell sorter (FACS) buffer (D-PBS-2% FCS) and then incubated with a saturating amount of the primary mouse MAb for 45 min on ice. After two washes, cells were incubated with PE-labeled goat anti-mouse Ig for 30 min on ice. Complexes of NCR-Fc fusion proteins (1 to 2 μg per staining) and PE-labeled goat anti-hIgG Fc (Dianova; 1:100 in FACS buffer) were allowed to form for 30 min before being added to cells for 60 min on ice. Cells were washed twice and re-suspended in 200 μl of FACS buffer with 0.05% propidium iodide (Sigma-Aldrich). Cytofluorometric analyses were done using a Canto2 (three lasers) or Fortesa3 (four lasers) flow cytometer and Diva software (Becton Dickinson, Heidelberg, Germany). For all FACS staining’s, representative examples are shown from at least three repeats with similar results.

### P24 antigen capture assay

Lentivirus particles based on the human immunodeficiency virus-1 (HIV-1) were produced in HEK-293 cells through transient transfection of 2 plasmids encoding components of the virus envelope. Cell culture medium containing viral particles produced by packaging cells was harvested after 72 h. HEK-293 cells stably transfected with the plasmid pcDNA6/V5 HisB-EBOV-GP) and control HEK-293 cells stably transfected with pcDNA6/V5 HisB were used to produce lentiviral particles displaying EBOV-GP (lenti-EBOV-GP) and control particles devoid of EBOV-GP. Briefly, 1×10^6^ cells of each cell line were seeded in 60 mm cell culture dishes one night prior to transfection. The next day, 6μg of a 3: 2 ratio of lentiviral transfer vector (pLOX-CWgfp) and packaging plasmid (psPAX2) were transfected into each cell culture dish using Turbofect Transfection Reagent (Thermo Fisher Scientific). 72 h post-transfection, viral supernatants were collected and centrifuged at 3000×g for 15 min at 4°C to remove cell debris. To concentrate the viral particles, they were centrifuged at 48,000×g for 3 h at 4°C, and viral pellets were re-suspended in cold PBS. Physical titration of the viral preparation was performed using p24 detection by sandwich ELISA. Microtiter plates pre-coated with anti-p24 Ab were incubated with increasing dilutions of the lentiviral suspension. After incubation and washing, p24 was quantified using a biotinylated anti-p24 Ab and detected using HRP-streptavidin. Color development was measured at 450 nm in a Bio-Rad spectrophotometer.

### ELISA Assay (lenti-EBOV-GP)

Lentiviral vector titers are expressed in transducing units per ml. After physical titration of viral vectors, we immobilized HIV-lentiviral EBOV-GP particles (200 μg/ml) on 96-well microtiter plates (Nunc, Maxisorp) in 0.1 M sodium bicarbonate buffer (pH 9.6) at 4°C for 18h and blocked with protein-free blocking buffer (PBS) pH 7.4 (Thermo), which is recommended for viral particles with highly glycosylated proteins (183). For the coating with EBOV-GP and control lentiviral particles, equal amounts of particles were added to each well. After blocking, the plates were washed three times with PBS containing 0.05% Tween-20. Dilutions of 10 μg/ml of fusion proteins were prepared in PBS + 2% protein free buffer, including NKp44-Fc, NKp46-Fc, NKp30-Fc, DC-SIGN-Fc, DC-SIGNR-Fc, Siglec2-Fc, Siglec3-Fc, Siglec4-Fc, Siglec5-Fc, Siglec7-Fc, Siglec10-Fc, PSGL-1-Fc,NKG2D-Fc, E-selectin-Fc, P-selectin-Fc, L-selectin-Fc, TIM1-Fc, CD44-Fc, CD24-Fc, syndecan 1-Fc (R&D) and human anti-Gp-Fc antibody as positive control. Each protein was added to the respective wells and incubated for 1h at room temperature. After incubation, the plates were gently washed three times with PBS containing 0.05% Tween-20. Then, the secondary antibody (goat anti-human IgG-Fc) was added to the wells and the plates incubated for 1h at room temperature. The plates were subsequently washed three times with PBS-T (PBS containing 0.05% Tween-20) and incubated with substrate (OPD) for 15-20 min. The reactions were stopped by adding 100μl of 1M sulfuric acid to each well. Finally, the absorbance of 450/570 nm was measured by a BioTek Synergy 4 Multi-Mode Microplate Reader.

## Acknowledgements

We are thankful to Professor Stephan Becker (Institute of Virology, of the Philipps-University, Marburg) for kindly providing the HEK-293 EVOB transfectant and the EVOB-GP plasmid.

## Footnotes

1 Syndecan receptors are required for internalization of the HIV-1 tat protein.

2 (Celularity CEO Robert Hariri; Xiaokui Zhang, Ph.D. “CYNK-001 demonstrates a range of biological activities expected of NK cells, including expression of activating receptors such as NKG2D, DNAM-1 and the natural cytotoxicity receptors NKp30, NKp44 and NKp46, which bind to stress ligands and viral antigens on infected cells. They also show the expression of cytolytic molecules perforin and granzyme B, which kill recognized infected cells. These functions suggest that CYNK-001 could provide a benefit to COVID-19 patients in terms of limiting SARS-CoV-2 replication and disease progression by eliminating the infected cells.” https://www.fiercebiotech.com/biotech/rudy-giuliani-backed-covid-19-therapy-from-celularity-nabs-fda-speedy-trial-start)

3 The CHO-K1 cell line (ATCC CCL-61) was derived as a subclone from the parental CHO cell line initiated from a biopsy of an ovary of an adult Chinese hamster by T. T. Puck in 1957. The cells require proline in the medium for growth.

## References

1. Bray M, Mahanty S. Ebola hemorrhagic fever and septic shock. J Infect Dis. 2003;188(11):1613–7.

2. Feldmann H, Geisbert TW. Ebola haemorrhagic fever. Lancet. 2011;377(9768):849–62.

3. Bray M, Murphy FA. Filovirus research: knowledge expands to meet a growing threat. J Infect Dis. 2007;196 Suppl 2:S438–43.

4. Kortepeter MG, Bausch DG, Bray M. Basic clinical and laboratory features of filoviral hemorrhagic fever. J Infect Dis. 2011;204 Suppl 3:S810–6.

5. Cardenas WB, Loo YM, Gale M, Jr., Hartman AL, Kimberlin CR, Martinez-Sobrido L, et al. Ebola virus VP35 protein binds double-stranded RNA and inhibits alpha/beta interferon production induced by RIG-I signaling. J Virol. 2006;80(11):5168–78.

6. Keshtkar-Jahromi M, Martins KAO, Cardile AP, Reisler RB, Christopher GW, Bavari S. Treatment-focused Ebola trials, supportive care and future of filovirus care. Expert Rev Anti Infect Ther. 2018;16(1):67–76.

7. Licata JM, Johnson RF, Han Z, Harty RN. Contribution of ebola virus glycoprotein, nucleoprotein, and VP24 to budding of VP40 virus-like particles. J Virol. 2004;78(14):7344–51.

8. Volchkov VE, Feldmann H, Volchkova VA, Klenk HD. Processing of the Ebola virus glycoprotein by the proprotein convertase furin. Proc Natl Acad Sci U S A. 1998;95(10):5762–7.

9. Barrientos LG, Martin AM, Rollin PE, Sanchez A. Disulfide bond assignment of the Ebola virus secreted glycoprotein SGP. Biochem Biophys Res Commun. 2004;323(2):696–702.

10. Falzarano D, Krokhin O, Wahl-Jensen V, Seebach J, Wolf K, Schnittler HJ, et al. Structure-function analysis of the soluble glycoprotein, sGP, of Ebola virus. Chembiochem. 2006;7(10):1605–11.

11. Sanchez A, Yang ZY, Xu L, Nabel GJ, Crews T, Peters CJ. Biochemical analysis of the secreted and virion glycoproteins of Ebola virus. J Virol. 1998;72(8):6442–7.

12. Stroher U, West E, Bugany H, Klenk HD, Schnittler HJ, Feldmann H. Infection and activation of monocytes by Marburg and Ebola viruses. J Virol. 2001;75(22):11025–33.

13. Klenk HD, Feldmann H. Symposium on Marburg and Ebola viruses. Virus Res. 2001;80(1-2):117–23.

14. Trefry JC, Wollen SE, Nasar F, Shamblin JD, Kern SJ, Bearss JJ, et al. Ebola Virus Infections in Nonhuman Primates Are Temporally Influenced by Glycoprotein Poly-U Editing Site Populations in the Exposure Material. Viruses. 2015;7(12):6739–54.

15. Jeffers SA, Sanders DA, Sanchez A. Covalent modifications of the ebola virus glycoprotein. J Virol. 2002;76(24):12463–72.

16. Volchkov VE, Volchkova VA, Slenczka W, Klenk HD, Feldmann H. Release of viral glycoproteins during Ebola virus infection. Virology. 1998;245(1):110–9.

17. Iwasa A, Shimojima M, Kawaoka Y. sGP serves as a structural protein in Ebola virus infection. J Infect Dis. 2011;204 Suppl 3:S897–903.

18. Feldmann F, Feldmann H. Ebola: facing a new transboundary animal disease? Dev Biol (Basel). 2013;135:201–9.

19. Miller EH, Obernosterer G, Raaben M, Herbert AS, Deffieu MS, Krishnan A, et al. Ebola virus entry requires the host-programmed recognition of an intracellular receptor. EMBO J. 2012;31(8):1947–60.

20. Beniac DR, Booth TF. Structure of the Ebola virus glycoprotein spike within the virion envelope at 11 A resolution. Sci Rep. 2017;7:46374.

21. Ray RB, Basu A, Steele R, Beyene A, McHowat J, Meyer K, et al. Ebola virus glycoprotein-mediated anoikis of primary human cardiac microvascular endothelial cells. Virology. 2004;321(2):181–8.

22. Simmons G, Wool-Lewis RJ, Baribaud F, Netter RC, Bates P. Ebola virus glycoproteins induce global surface protein down-modulation and loss of cell adherence. J Virol. 2002;76(5):2518–28.

23. Takada A, Watanabe S, Ito H, Okazaki K, Kida H, Kawaoka Y. Downregulation of beta1 integrins by Ebola virus glycoprotein: implication for virus entry. Virology. 2000;278(1):20–6.

24. Yang ZY, Duckers HJ, Sullivan NJ, Sanchez A, Nabel EG, Nabel GJ. Identification of the Ebola virus glycoprotein as the main viral determinant of vascular cell cytotoxicity and injury. Nat Med. 2000;6(8):886–9.

25. Zhang X, Wang C, Chen B, Wang Q, Xu W, Ye S, et al. Crystal Structure of Refolding Fusion Core of Lassa Virus GP2 and Design of Lassa Virus Fusion Inhibitors. Front Microbiol. 2019;10:1829.

26. Hastie KM, Zandonatti MA, Kleinfelter LM, Heinrich ML, Rowland MM, Chandran K, et al. Structural basis for antibody-mediated neutralization of Lassa virus. Science. 2017;356(6341):923–8.

27. Wahl-Jensen V, Kurz SK, Hazelton PR, Schnittler HJ, Stroher U, Burton DR, et al. Role of Ebola virus secreted glycoproteins and virus-like particles in activation of human macrophages. Journal of Virology. 2005;79(4):2413–9.

28. Wahl-Jensen VM, Afanasieva TA, Seebach J, Stroher U, Feldmann H, Schnittler HJ. Effects of Ebola virus glycoproteins on endothelial cell activation and barrier function. Journal of Virology. 2005;79(16):10442–50.

29. Falzarano D, Krokhin O, Van Domselaar G, Wolf K, Seebach J, Schnittler HJ, et al. Ebola sGP--the first viral glycoprotein shown to be C-mannosylated. Virology. 2007;368(1):83–90.

30. Fuller CL, Ruthel G, Warfield KL, Swenson DL, Bosio CM, Aman MJ, et al. NKp30-dependent cytolysis of filovirus-infected human dendritic cells. Cell Microbiol. 2007;9(4):962–76.

31. Galandrini R, Palmieri G, Piccoli M, Frati L, Santoni A. Role for the Rac1 exchange factor Vav in the signaling pathways leading to NK cell cytotoxicity. J Immunol. 1999;162(6):3148–52.

32. Villalba M, Hernandez J, Deckert M, Tanaka Y, Altman A. Vav modulation of the Ras/MEK/ERK signaling pathway plays a role in NFAT activation and CD69 up-regulation. Eur J Immunol. 2000;30(6):1587–96.

33. Ansari AA. Clinical features and pathobiology of Ebolavirus infection. J Autoimmun. 2014;55:1–9.

34. Misasi J, Sullivan NJ. Camouflage and misdirection: the full-on assault of ebola virus disease. Cell. 2014;159(3):477–86.

35. Bosio CM, Aman MJ, Grogan C, Hogan R, Ruthel G, Negley D, et al. Ebola and Marburg viruses replicate in monocyte-derived dendritic cells without inducing the production of cytokines and full maturation. J Infect Dis. 2003;188(11):1630–8.

36. Melanson VR, Kalina WV, Williams P. Ebola virus infection induces irregular dendritic cell gene expression. Viral Immunol. 2015;28(1):42–50.

37. Mahanty S, Gupta M, Paragas J, Bray M, Ahmed R, Rollin PE. Protection from lethal infection is determined by innate immune responses in a mouse model of Ebola virus infection. Virology. 2003;312(2):415–24.

38. Mahanty S, Hutchinson K, Agarwal S, McRae M, Rollin PE, Pulendran B. Cutting edge: impairment of dendritic cells and adaptive immunity by Ebola and Lassa viruses. J Immunol. 2003;170(6):2797–801.

39. Moretta L, Moretta A. Unravelling natural killer cell function: triggering and inhibitory human NK receptors. EMBO J. 2004;23(2):255–9.

40. Sivori S, Vitale M, Morelli L, Sanseverino L, Augugliaro R, Bottino C, et al. p46, a novel natural killer cell-specific surface molecule that mediates cell activation. J Exp Med. 1997;186(7):1129–36.

41. Pessino A, Sivori S, Bottino C, Malaspina A, Morelli L, Moretta L, et al. Molecular cloning of NKp46: a novel member of the immunoglobulin superfamily involved in triggering of natural cytotoxicity. J Exp Med. 1998;188(5):953–60.

42. Verrier T, Satoh-Takayama N, Serafini N, Marie S, Di Santo JP, Vosshenrich CA. Phenotypic and Functional Plasticity of Murine Intestinal NKp46+ Group 3 Innate Lymphoid Cells. J Immunol. 2016;196(11):4731–8.

43. Correia MP, Costa AV, Uhrberg M, Cardoso EM, Arosa FA. IL-15 induces CD8+ T cells to acquire functional NK receptors capable of modulating cytotoxicity and cytokine secretion. Immunobiology. 2011;216(5):604–12.

44. Vitale M, Bottino C, Sivori S, Sanseverino L, Castriconi R, Marcenaro E, et al. NKp44, a novel triggering surface molecule specifically expressed by activated natural killer cells, is involved in non-major histocompatibility complex-restricted tumor cell lysis. J Exp Med. 1998;187(12):2065–72.

45. Cantoni C, Bottino C, Vitale M, Pessino A, Augugliaro R, Malaspina A, et al. NKp44, a triggering receptor involved in tumor cell lysis by activated human natural killer cells, is a novel member of the immunoglobulin superfamily. J Exp Med. 1999;189(5):787–96.

46. Shemesh A, Kugel A, Steiner N, Yezersky M, Tirosh D, Edri A, et al. NKp44 and NKp30 splice variant profiles in decidua and tumor tissues: a comparative viewpoint. Oncotarget. 2016;7(43):70912–23.

47. Lanier LL. Activating and inhibitory NK cell receptors. Adv Exp Med Biol. 1998;452:13–8.

48. Cantoni C, Ponassi M, Biassoni R, Conte R, Spallarossa A, Moretta A, et al. The three-dimensional structure of the human NK cell receptor NKp44, a triggering partner in natural cytotoxicity. Structure. 2003;11(6):725–34.

49. Campbell KS, Yusa S, Kikuchi-Maki A, Catina TL. NKp44 triggers NK cell activation through DAP12 association that is not influenced by a putative cytoplasmic inhibitory sequence. J Immunol. 2004;172(2):899–906.

50. Salimi M, Xue LZ, John H, Hardman C, Cousins DJ, McKenzie ANJ, et al. Group 2 Innate Lymphoid Cells Express Functional NKp30 Receptor Inducing Type 2 Cytokine Production. Journal of Immunology. 2016;196(1):45–54.

51. Pende D, Parolini S, Pessino A, Sivori S, Augugliaro R, Morelli L, et al. Identification and molecular characterization of NKp30, a novel triggering receptor involved in natural cytotoxicity mediated by human natural killer cells. J Exp Med. 1999;190(10):1505–16.

52. Moretta A, Bottino C, Vitale M, Pende D, Cantoni C, Mingari MC, et al. Activating receptors and coreceptors involved in human natural killer cell-mediated cytolysis. Annu Rev Immunol. 2001;19:197–223.

53. Schlums H, Cichocki F, Tesi B, Theorell J, Beziat V, Holmes TD, et al. Cytomegalovirus infection drives adaptive epigenetic diversification of NK cells with altered signaling and effector function. Immunity. 2015;42(3):443–56.

54. Kaifu T, Escaliere B, Gastinel LN, Vivier E, Baratin M. B7-H6/NKp30 interaction: a mechanism of alerting NK cells against tumors. Cell Mol Life Sci. 2011;68(21):3531–9.

55. Brandt CS, Baratin M, Yi EC, Kennedy J, Gao Z, Fox B, et al. The B7 family member B7-H6 is a tumor cell ligand for the activating natural killer cell receptor NKp30 in humans. J Exp Med. 2009;206(7):1495–503.

56. Byrd A, Hoffmann SC, Jarahian M, Momburg F, Watzl C. Expression Analysis of the Ligands for the Natural Killer Cell Receptors NKp30 and NKp44. Plos One. 2007;2(12).

57. Hecht ML, Rosental B, Horlacher T, Hershkovitz O, De Paz JL, Noti C, et al. Natural cytotoxicity receptors NKp30, NKp44 and NKp46 bind to different heparan sulfate/heparin sequences. J Proteome Res. 2009;8(2):712–20.

58. Hershkovitz O, Jarahian M, Zilka A, Bar-Ilan A, Landau G, Jivov S, et al. Altered glycosylation of recombinant NKp30 hampers binding to heparan sulfate: a lesson for the use of recombinant immunoreceptors as an immunological tool. Glycobiology. 2008;18(1):28–41.

59. Jarahian M, Fiedler M, Cohnen A, Djandji D, Hammerling GJ, Gati C, et al. Modulation of NKp30- and NKp46-mediated natural killer cell responses by poxviral hemagglutinin. PLoS Pathog. 2011;7(8):e1002195.

60. Arnon TI, Achdout H, Levi O, Markel G, Saleh N, Katz G, et al. Inhibition of the NKp30 activating receptor by pp65 of human cytomegalovirus. Nat Immunol. 2005;6(5):515–23.

61. Delahaye NF, Rusakiewicz S, Martins I, Menard C, Roux S, Lyonnet L, et al. Alternatively spliced NKp30 isoforms affect the prognosis of gastrointestinal stromal tumors. Nat Med. 2011;17(6):700–7.

62. Karupiah G, Blanden RV, Ramshaw IA. Interferon gamma is involved in the recovery of athymic nude mice from recombinant vaccinia virus/interleukin 2 infection. J Exp Med. 1990;172(5):1495–503.

63. Parker AK, Parker S, Yokoyama WM, Corbett JA, Buller RM. Induction of natural killer cell responses by ectromelia virus controls infection. J Virol. 2007;81(8):4070–9.

64. Martinez J, Huang X, Yang Y. Direct action of type I IFN on NK cells is required for their activation in response to vaccinia viral infection in vivo. J Immunol. 2008;180(3):1592–7.

65. Chisholm SE, Reyburn HT. Recognition of vaccinia virus-infected cells by human natural killer cells depends on natural cytotoxicity receptors. J Virol. 2006;80(5):2225–33.

66. Chisholm SE, Howard K, Gomez MV, Reyburn HT. Expression of ICP0 is sufficient to trigger natural killer cell recognition of herpes simplex virus-infected cells by natural cytotoxicity receptors. J Infect Dis. 2007;195(8):1160–8.

67. Vieillard V, Strominger JL, Debre P. NK cytotoxicity against CD4+ T cells during HIV-1 infection: a gp41 peptide induces the expression of an NKp44 ligand. Proc Natl Acad Sci U S A. 2005;102(31):10981–6.

68. Bar-On Y, Charpak-Amikam Y, Glasner A, Isaacson B, Duev-Cohen A, Tsukerman P, et al. NKp46 Recognizes the Sigma1 Protein of Reovirus: Implications for Reovirus-Based Cancer Therapy. J Virol. 2017;91(19).

69. Mandelboim O, Lieberman N, Lev M, Paul L, Arnon TI, Bushkin Y, et al. Recognition of haemagglutinins on virus-infected cells by NKp46 activates lysis by human NK cells. Nature. 2001;409(6823):1055–60.

70. Gazit R, Gruda R, Elboim M, Arnon TI, Katz G, Achdout H, et al. Lethal influenza infection in the absence of the natural killer cell receptor gene Ncr1. Nat Immunol. 2006;7(5):517–23.

71. Arnon TI, Lev M, Katz G, Chernobrov Y, Porgador A, Mandelboim O. Recognition of viral hemagglutinins by NKp44 but not by NKp30. Eur J Immunol. 2001;31(9):2680–9.

72. Jarahian M, Watzl C, Fournier P, Arnold A, Djandji D, Zahedi S, et al. Activation of natural killer cells by newcastle disease virus hemagglutinin-neuraminidase. J Virol. 2009;83(16):8108–21.

73. Lenman A, Liaci AM, Liu Y, Frangsmyr L, Frank M, Blaum BS, et al. Polysialic acid is a cellular receptor for human adenovirus 52. Proc Natl Acad Sci U S A. 2018;115(18):E4264–E73.

74. Hershkovitz O, Rosental B, Rosenberg LA, Navarro-Sanchez ME, Jivov S, Zilka A, et al. NKp44 receptor mediates interaction of the envelope glycoproteins from the West Nile and dengue viruses with NK cells. J Immunol. 2009;183(4):2610–21.

75. Gramberg T, Soilleux E, Fisch T, Lalor PF, Hofmann H, Wheeldon S, et al. Interactions of LSECtin and DC-SIGN/DC-SIGNR with viral ligands: Differential pH dependence, internalization and virion binding. Virology. 2008;373(1):189–201.

76. Leger P, Tetard M, Youness B, Cordes N, Rouxel RN, Flamand M, et al. Differential Use of the C-Type Lectins L-SIGN and DC-SIGN for Phlebovirus Endocytosis. Traffic. 2016;17(6):639–56.

77. Lozach PY, Amara A, Bartosch B, Virelizier JL, Arenzana-Seisdedos F, Cosset FL, et al. C-type lectins L-SIGN and DC-SIGN capture and transmit infectious hepatitis C virus pseudotype particles. J Biol Chem. 2004;279(31):32035–45.

78. Lozach PY, Kuhbacher A, Meier R, Mancini R, Bitto D, Bouloy M, et al. DC-SIGN as a receptor for phleboviruses. Cell Host Microbe. 2011;10(1):75–88.

79. Khoo US, Chan KYK, Chan VSF, Lin CLS. DC-SIGN and L-SIGN: the SIGNs for infection. J Mol Med. 2008;86(8):861–74.

80. Nabatov AA, Raginov IS. The DC-SIGN-CD56 interaction inhibits the anti-dendritic cell cytotoxicity of CD56 expressing cells. Infect Agent Cancer. 2015;10:49.

81. Goncalves AR, Moraz ML, Pasquato A, Helenius A, Lozach PY, Kunz S. Role of DC-SIGN in Lassa virus entry into human dendritic cells. J Virol. 2013;87(21):11504–15.

82. Maier I, Wu GY. Hepatitis C and HIV co-infection: a review. World J Gastroentero. 2002;8(4):577–9.

83. Raffray L, Giry C, Thirapathi Y, Reboux AH, Jaffar-Bandjee MC, Gasque P. Increased levels of soluble forms of E-selectin and ICAM-1 adhesion molecules during human leptospirosis. PLoS One. 2017;12(7):e0180474.

84. Vassena L, Giuliani E, Koppensteiner H, Bolduan S, Schindler M, Doria M. HIV-1 Nef and Vpu Interfere with L-Selectin (CD62L) Cell Surface Expression To Inhibit Adhesion and Signaling in Infected CD4+ T Lymphocytes. J Virol. 2015;89(10):5687–700.

85. Kononchik J, Ireland J, Zou Z, Segura J, Holzapfel G, Chastain A, et al. HIV-1 targets L-selectin for adhesion and induces its shedding for viral release. Nat Commun. 2018;9(1):2825.

86. Ivetic A, Hoskins Green HL, Hart SJ. L-selectin: A Major Regulator of Leukocyte Adhesion, Migration and Signaling. Front Immunol. 2019;10:1068.

87. Hernandez Mir G, Helin J, Skarp KP, Cummings RD, Makitie A, Renkonen R, et al. Glycoforms of human endothelial CD34 that bind L-selectin carry sulfated sialyl Lewis x capped O- and N-glycans. Blood. 2009;114(3):733–41.

88. Kondratowicz AS, Lennemann NJ, Sinn PL, Davey RA, Hunt CL, Moller-Tank S, et al. T-cell immunoglobulin and mucin domain 1 (TIM-1) is a receptor for Zaire Ebolavirus and Lake Victoria Marburgvirus. Proc Natl Acad Sci U S A. 2011;108(20):8426–31.

89. Jemielity S, Wang JJ, Chan YK, Ahmed AA, Li W, Monahan S, et al. TIM-family proteins promote infection of multiple enveloped viruses through virion-associated phosphatidylserine. PLoS Pathog. 2013;9(3):e1003232.

90. Moller-Tank S, Kondratowicz AS, Davey RA, Rennert PD, Maury W. Role of the phosphatidylserine receptor TIM-1 in enveloped-virus entry. J Virol. 2013;87(15):8327–41.

91. Morizono K, Chen IS. Role of phosphatidylserine receptors in enveloped virus infection. J Virol. 2014;88(8):4275–90.

92. Angiari S, Constantin G. Regulation of T cell trafficking by the T cell immunoglobulin and mucin domain 1 glycoprotein. Trends Mol Med. 2014;20(12):675–84.

93. Angiari S, Donnarumma T, Rossi B, Dusi S, Pietronigro E, Zenaro E, et al. TIM-1 glycoprotein binds the adhesion receptor P-selectin and mediates T cell trafficking during inflammation and autoimmunity. Immunity. 2014;40(4):542–53.

94. Du P, Xiong R, Li X, Jiang J. Immune Regulation and Antitumor Effect of TIM-1. J Immunol Res. 2016;2016:8605134.

95. Graham DK, DeRyckere D, Davies KD, Earp HS. The TAM family: phosphatidylserine sensing receptor tyrosine kinases gone awry in cancer. Nat Rev Cancer. 2014;14(12):769–85.

96. Rhein BA, Maury WJ. Ebola virus entry into host cells: identifying therapeutic strategies. Curr Clin Microbiol Rep. 2015;2(3):115–24.

97. Moller-Tank S, Maury W. Ebola virus entry: a curious and complex series of events. PLoS Pathog. 2015;11(4):e1004731.

98. van der Meer JH, van der Poll T, van’t Veer C. TAM receptors, Gas6, and protein S: roles in inflammation and hemostasis. Blood. 2014;123(16):2460–9.

99. Falasca L, Agrati C, Petrosillo N, Di Caro A, Capobianchi MR, Ippolito G, et al. Molecular mechanisms of Ebola virus pathogenesis: focus on cell death. Cell Death Differ. 2015;22(8):1250–9.

100. Chan SY, Speck RF, Ma MC, Goldsmith MA. Distinct mechanisms of entry by envelope glycoproteins of Marburg and Ebola (Zaire) viruses. J Virol. 2000;74(10):4933–7.

101. Lee JE, Fusco ML, Hessell AJ, Oswald WB, Burton DR, Saphire EO. Structure of the Ebola virus glycoprotein bound to an antibody from a human survivor. Nature. 2008;454(7201):177–82.

102. Lucht A, Grunow R, Otterbein C, Moller P, Feldmann H, Becker S. Production of monoclonal antibodies and development of an antigen capture ELISA directed against the envelope glycoprotein GP of Ebola virus. Med Microbiol Immunol. 2004;193(4):181–7.

103. Zhong X, Ma W, Meade CL, Tam AS, Llewellyn E, Cornell R, et al. Transient CHO expression platform for robust antibody production and its enhanced N-glycan sialylation on therapeutic glycoproteins. Biotechnol Prog. 2019;35(1):e2724.

104. Campbell C, Stanley P. A dominant mutation to ricin resistance in Chinese hamster ovary cells induces UDP-GlcNAc:glycopeptide beta-4-N-acetylglucosaminyltransferase III activity. J Biol Chem. 1984;259(21):13370–8.

105. Stanley P, Sundaram S, Tang J, Shi S. Molecular analysis of three gain-of-function CHO mutants that add the bisecting GlcNAc to N-glycans. Glycobiology. 2005;15(1):43–53.

106. Harduin-Lepers A, Vallejo-Ruiz V, Krzewinski-Recchi MA, Samyn-Petit B, Julien S, Delannoy P. The human sialyltransferase family. Biochimie. 2001;83(8):727–37.

107. Li SS, Ip CKM, Tang MYH, Tang MKS, Tong Y, Zhang J, et al. Sialyl Lewis(x)-P-selectin cascade mediates tumor-mesothelial adhesion in ascitic fluid shear flow. Nat Commun. 2019;10(1):2406.

108. Dimitroff CJ, Lee JY, Rafii S, Fuhlbrigge RC, Sackstein R. CD44 is a major E-selectin ligand on human hematopoietic progenitor cells. J Cell Biol. 2001;153(6):1277–86.

109. Gout S, Morin C, Houle F, Huot J. Death receptor-3, a new E-Selectin counter-receptor that confers migration and survival advantages to colon carcinoma cells by triggering p38 and ERK MAPK activation. Cancer Res. 2006;66(18):9117–24.

110. Kitamura K, Sato K, Sawabe M, Yoshida M, Hagiwara N. P-Selectin Glycoprotein Ligand-1 (PSGL-1) Expressing CD4 T Cells Contribute Plaque Instability in Acute Coronary Syndrome. Circ J. 2018;82(8):2128–35.

111. Kappelmayer J, Nagy B, Jr. The Interaction of Selectins and PSGL-1 as a Key Component in Thrombus Formation and Cancer Progression. Biomed Res Int. 2017;2017:6138145.

112. Park H, Kim M, Kim HJ, Lee Y, Seo Y, Pham CD, et al. Heparan sulfate proteoglycans (HSPGs) and chondroitin sulfate proteoglycans (CSPGs) function as endocytic receptors for an internalizing anti-nucleic acid antibody. Sci Rep. 2017;7(1):14373.

113. Birzele F, Voss E, Nopora A, Honold K, Heil F, Lohmann S, et al. CD44 Isoform Status Predicts Response to Treatment with Anti-CD44 Antibody in Cancer Patients. Clin Cancer Res. 2015;21(12):2753–62.

114. Barthel SR, Gavino JD, Wiese GK, Jaynes JM, Siddiqui J, Dimitroff CJ. Analysis of glycosyltransferase expression in metastatic prostate cancer cells capable of rolling activity on microvascular endothelial (E)-selectin. Glycobiology. 2008;18(10):806–17.

115. Edri A, Shemesh A, Iraqi M, Matalon O, Brusilovsky M, Hadad U, et al. The Ebola-Glycoprotein Modulates the Function of Natural Killer Cells. Front Immunol. 2018;9:1428.

116. Sanders WJ, Katsumoto TR, Bertozzi CR, Rosen SD, Kiessling LL. L-selectin-carbohydrate interactions: relevant modifications of the Lewis x trisaccharide. Biochemistry. 1996;35(47):14862–7.

117. Sperandio M, Gleissner CA, Ley K. Glycosylation in immune cell trafficking. Immunol Rev. 2009;230(1):97–113.

118. de Leeuw O, Peeters B. Complete nucleotide sequence of Newcastle disease virus: evidence for the existence of a new genus within the subfamily Paramyxovirinae. J Gen Virol. 1999;80 (Pt 1):131–6.

119. Tremblay MJ, Fortin JF, Cantin R. The acquisition of host-encoded proteins by nascent HIV-1. Immunol Today. 1998;19(8):346–51.

120. Segura MM, Garnier A, Di Falco MR, Whissell G, Meneses-Acosta A, Arcand N, et al. Identification of host proteins associated with retroviral vector particles by proteomic analysis of highly purified vector preparations. J Virol. 2008;82(3):1107–17.

121. Giguere JF, Paquette JS, Bounou S, Cantin R, Tremblay MJ. New insights into the functionality of a virion-anchored host cell membrane protein: CD28 versus HIV type 1. J Immunol. 2002;169(5):2762–71.

122. Louder MK, Sambor A, Chertova E, Hunte T, Barrett S, Ojong F, et al. HIV-1 envelope pseudotyped viral vectors and infectious molecular clones expressing the same envelope glycoprotein have a similar neutralization phenotype, but culture in peripheral blood mononuclear cells is associated with decreased neutralization sensitivity. Virology. 2005;339(2):226–38.

123. Bloushtain N, Qimron U, Bar-Ilan A, Hershkovitz O, Gazit R, Fima E, et al. Membrane-associated heparan sulfate proteoglycans are involved in the recognition of cellular targets by NKp30 and NKp46. J Immunol. 2004;173(4):2392–401.

124. Giroglou T, Florin L, Schafer F, Streeck RE, Sapp M. Human papillomavirus infection requires cell surface heparan sulfate. J Virol. 2001;75(3):1565–70.

125. Shafti-Keramat S, Handisurya A, Kriehuber E, Meneguzzi G, Slupetzky K, Kirnbauer R. Different heparan sulfate proteoglycans serve as cellular receptors for human papillomaviruses. J Virol. 2003;77(24):13125–35.

126. Lanitis E, Dangaj D, Irving M, Coukos G. Mechanisms regulating T-cell infiltration and activity in solid tumors. Ann Oncol. 2017;28(suppl_12):xii18–xii32.

127. Gajewski TF, Schreiber H, Fu YX. Innate and adaptive immune cells in the tumor microenvironment. Nat Immunol. 2013;14(10):1014–22.

128. Moretta A, Bottino C, Sivori S, Marcenaro E, Castriconi R, Della Chiesa M, et al. Natural killer lymphocytes: “null cells” no more. Ital J Anat Embryol. 2001;106(4):335–42.

129. Moretta L, Biassoni R, Bottino C, Mingari MC, Moretta A. Immunobiology of human NK cells. Transplant Proc. 2001;33(1-2):60–1.

130. De Pelsmaeker S, Romero N, Vitale M, Favoreel HW. Herpesvirus Evasion of Natural Killer Cells. J Virol. 2018;92(11).

131. De Pelsmaeker S, Devriendt B, De Regge N, Favoreel HW. Porcine NK Cells Stimulate Proliferation of Pseudorabies Virus-Experienced CD8(+) and CD4(+)CD8(+) T Cells. Front Immunol. 2018;9:3188.

132. Chiang SCC, Wood SM, Tesi B, Akar HH, Al-Herz W, Ammann S, et al. Differences in Granule Morphology yet Equally Impaired Exocytosis among Cytotoxic T Cells and NK Cells from Chediak-Higashi Syndrome Patients. Front Immunol. 2017;8:426.

133. Krzewski K, Gil-Krzewska A, Nguyen V, Peruzzi G, Coligan JE. LAMP1/CD107a is required for efficient perforin delivery to lytic granules and NK-cell cytotoxicity. Blood. 2013;121(23):4672–83.

134. Krzewski K, Coligan JE. Human NK cell lytic granules and regulation of their exocytosis. Front Immunol. 2012;3:335.

135. Long EO, Rajagopalan S. Stress signals activate natural killer cells. J Exp Med. 2002;196(11):1399–402.

136. Zamai L, Ahmad M, Bennett IM, Azzoni L, Alnemri ES, Perussia B. Natural killer (NK) cell-mediated cytotoxicity: differential use of TRAIL and Fas ligand by immature and mature primary human NK cells. J Exp Med. 1998;188(12):2375–80.

137. Boelen L, Debebe B, Silveira M, Salam A, Makinde J, Roberts CH, et al. Inhibitory killer cell immunoglobulin-like receptors strengthen CD8(+) T cell-mediated control of HIV-1, HCV, and HTLV-1. Sci Immunol. 2018;3(29).

138. Ploquin A, Zhou Y, Sullivan NJ. Ebola Immunity: Gaining a Winning Position in Lightning Chess. J Immunol. 2018;201(3):833–42.

139. Younan P, Iampietro M, Bukreyev A. Disabling of lymphocyte immune response by Ebola virus. PLoS Pathog. 2018;14(4):e1006932.

140. Warfield KL, Perkins JG, Swenson DL, Deal EM, Bosio CM, Aman MJ, et al. Role of natural killer cells in innate protection against lethal ebola virus infection. J Exp Med. 2004;200(2):169–79.

141. Fausther-Bovendo H, Qiu X, He S, Bello A, Audet J, Ippolito G, et al. NK Cells Accumulate in Infected Tissues and Contribute to Pathogenicity of Ebola Virus in Mice. J Virol. 2019;93(10).

142. Charpak-Amikam Y, Kubsch T, Seidel E, Oiknine-Djian E, Cavaletto N, Yamin R, et al. Human cytomegalovirus escapes immune recognition by NK cells through the downregulation of B7-H6 by the viral genes US18 and US20. Sci Rep. 2017;7(1):8661.

143. Alvarez CP, Lasala F, Carrillo J, Muniz O, Corbi AL, Delgado R. C-type lectins DC-SIGN and L-SIGN mediate cellular entry by Ebola virus in cis and in trans. J Virol. 2002;76(13):6841–4.

144. Simmons G, Reeves JD, Grogan CC, Vandenberghe LH, Baribaud F, Whitbeck JC, et al. DC-SIGN and DC-SIGNR bind ebola glycoproteins and enhance infection of macrophages and endothelial cells. Virology. 2003;305(1):115–23.

145. Takada A, Fujioka K, Tsuiji M, Morikawa A, Higashi N, Ebihara H, et al. Human macrophage C-type lectin specific for galactose and N-acetylgalactosamine promotes filovirus entry. J Virol. 2004;78(6):2943–7.

146. Li M, Ablan SD, Miao C, Zheng YM, Fuller MS, Rennert PD, et al. TIM-family proteins inhibit HIV-1 release. Proc Natl Acad Sci U S A. 2014;111(35):E3699–707.

147. Lee HH, Meyer EH, Goya S, Pichavant M, Kim HY, Bu X, et al. Apoptotic cells activate NKT cells through T cell Ig-like mucin-like-1 resulting in airway hyperreactivity. J Immunol. 2010;185(9):5225–35.

148. Prescher H, Frank M, Gutgemann S, Kuhfeldt E, Schweizer A, Nitschke L, et al. Design, Synthesis, and Biological Evaluation of Small, High-Affinity Siglec-7 Ligands: Toward Novel Inhibitors of Cancer Immune Evasion. J Med Chem. 2017;60(3):941–56.

149. Lucar O, Reeves RK, Jost S. A Natural Impact: NK Cells at the Intersection of Cancer and HIV Disease. Front Immunol. 2019;10:1850.

150. Zou F, Lu L, Liu J, Xia B, Zhang W, Hu Q, et al. Engineered triple inhibitory receptor resistance improves anti-tumor CAR-T cell performance via CD56. Nat Commun. 2019;10(1):4109.

151. Dragovich MA, Fortoul N, Jagota A, Zhang W, Schutt K, Xu Y, et al. Biomechanical characterization of TIM protein-mediated Ebola virus-host cell adhesion. Sci Rep. 2019;9(1):267.

152. Cognasse F, Nguyen KA, Damien P, McNicol A, Pozzetto B, Hamzeh-Cognasse H, et al. The Inflammatory Role of Platelets via Their TLRs and Siglec Receptors. Front Immunol. 2015;6:83.

153. Zhu W, Banadyga L, Emeterio K, Wong G, Qiu X. The Roles of Ebola Virus Soluble Glycoprotein in Replication, Pathogenesis, and Countermeasure Development. Viruses. 2019;11(11).

154. Bradfute SB, Warfield KL, Bavari S. Functional CD8+ T cell responses in lethal Ebola virus infection. J Immunol. 2008;180(6):4058–66.

155. Sanchez A, Lukwiya M, Bausch D, Mahanty S, Sanchez AJ, Wagoner KD, et al. Analysis of human peripheral blood samples from fatal and nonfatal cases of Ebola (Sudan) hemorrhagic fever: cellular responses, virus load, and nitric oxide levels. J Virol. 2004;78(19):10370–7.

156. Watzl C, Urlaub D, Fasbender F, Claus M. Natural killer cell regulation - beyond the receptors. F1000Prime Rep. 2014;6:87.

157. Beldi-Ferchiou A, Caillat-Zucman S. Control of NK Cell Activation by Immune Checkpoint Molecules. Int J Mol Sci. 2017;18(10).

158. Paul S, Lal G. The Molecular Mechanism of Natural Killer Cells Function and Its Importance in Cancer Immunotherapy. Front Immunol. 2017;8:1124.

159. Foley B, Felices M, Cichocki F, Cooley S, Verneris MR, Miller JS. The biology of NK cells and their receptors affects clinical outcomes after hematopoietic cell transplantation (HCT). Immunol Rev. 2014;258(1):45–63.

160. Avril T, Attrill H, Zhang J, Raper A, Crocker PR. Negative regulation of leucocyte functions by CD33-related siglecs. Biochem Soc T. 2006;34:1024–7.

161. Lubbers J, Rodriguez E, van Kooyk Y. Modulation of Immune Tolerance via Siglec-Sialic Acid Interactions. Frontiers in Immunology. 2018;9.

162. Stanczak MA, Siddiqui SS, Trefny MP, Thommen DS, Boligan KF, von Gunten S, et al. Self-associated molecular patterns mediate cancer immune evasion by engaging Siglecs on T cells. J Clin Invest. 2018;128(11):4912–23.

163. Cagnoni AJ, Perez Saez JM, Rabinovich GA, Marino KV. Turning-Off Signaling by Siglecs, Selectins, and Galectins: Chemical Inhibition of Glycan-Dependent Interactions in Cancer. Front Oncol. 2016;6:109.

164. Jandus C, Boligan KF, Chijioke O, Liu H, Dahlhaus M, Demoulins T, et al. Interactions between Siglec-7/9 receptors and ligands influence NK cell-dependent tumor immunosurveillance. J Clin Invest. 2014;124(4):1810–20.

165. Crocker PR, Paulson JC, Varki A. Siglecs and their roles in the immune system. Nat Rev Immunol. 2007;7(4):255–66.

166. Patel M, Yanagishita M, Roderiquez G, Bou-Habib DC, Oravecz T, Hascall VC, et al. Cell-surface heparan sulfate proteoglycan mediates HIV-1 infection of T-cell lines. AIDS Res Hum Retroviruses. 1993;9(2):167–74.

167. Xu Y, Martinez P, Seron K, Luo G, Allain F, Dubuisson J, et al. Characterization of hepatitis C virus interaction with heparan sulfate proteoglycans. J Virol. 2015;89(7):3846–58.

168. Kalia M, Chandra V, Rahman SA, Sehgal D, Jameel S. Heparan sulfate proteoglycans are required for cellular binding of the hepatitis E virus ORF2 capsid protein and for viral infection. J Virol. 2009;83(24):12714–24.

169. Salvador B, Sexton NR, Carrion R, Jr., Nunneley J, Patterson JL, Steffen I, et al. Filoviruses utilize glycosaminoglycans for their attachment to target cells. J Virol. 2013;87(6):3295–304.

170. O’Hearn A, Wang M, Cheng H, Lear-Rooney CM, Koning K, Rumschlag-Booms E, et al. Role of EXT1 and Glycosaminoglycans in the Early Stage of Filovirus Entry. J Virol. 2015;89(10):5441–9.

171. Lin CL, Chung CS, Heine HG, Chang W. Vaccinia virus envelope H3L protein binds to cell surface heparan sulfate and is important for intracellular mature virion morphogenesis and virus infection in vitro and in vivo. J Virol. 2000;74(7):3353–65.

172. Surviladze Z, Sterkand RT, Ozbun MA. Interaction of human papillomavirus type 16 particles with heparan sulfate and syndecan-1 molecules in the keratinocyte extracellular matrix plays an active role in infection. J Gen Virol. 2015;96(8):2232–41.

173. Whitley R, Kimberlin DW, Prober CG. Pathogenesis and disease. In: Arvin A, Campadelli-Fiume G, Mocarski E, Moore PS, Roizman B, Whitley R, et al., editors. Human Herpesviruses: Biology, Therapy, and Immunoprophylaxis. Cambridge2007.

174. Shukla D, Liu J, Blaiklock P, Shworak NW, Bai X, Esko JD, et al. A novel role for 3-O-sulfated heparan sulfate in herpes simplex virus 1 entry. Cell. 1999;99(1):13–22.

175. Tamhankar M, Gerhardt DM, Bennett RS, Murphy N, Jahrling PB, Patterson JL. Heparan sulfate is an important mediator of Ebola virus infection in polarized epithelial cells. Virol J. 2018;15(1):135.

176. Misra S, Heldin P, Hascall VC, Karamanos NK, Skandalis SS, Markwald RR, et al. Hyaluronan-CD44 interactions as potential targets for cancer therapy. Febs J. 2011;278(9):1429–43.

177. Ahrens T, Mai BH, Stingl AS, Termeer CC, Hofmann M, Fieber C, et al. Interaction of the extracellular matrix (ECM) component hyaluronan (HA) with its principle cell surface CD44 augments human melanoma cell proliferation. Journal of Investigative Dermatology. 1998;110(4):593-.

178. Lesley J, Hyman R, English N, Catterall JB, Turner GA. CD44 in inflammation and metastasis. Glycoconj J. 1997;14(5):611–22.

179. Guvench O. Revealing the Mechanisms of Protein Disorder and N-Glycosylation in CD44-Hyaluronan Binding Using Molecular Simulation. Front Immunol. 2015;6:305.

180. Nishimura Y, Shimojima M, Tano Y, Miyamura T, Wakita T, Shimizu H. Human P-selectin glycoprotein ligand-1 is a functional receptor for enterovirus 71. Nat Med. 2009;15(7):794–7.

181. Diab M, Glasner A, Isaacson B, Bar-On Y, Drori Y, Yamin R, et al. NK-cell receptors NKp46 and NCR1 control human metapneumovirus infection. Eur J Immunol. 2017;47(4):692–703.

182. Mandelboim O, Porgador A. NKp46. Int J Biochem Cell Biol. 2001;33(12):1147–50.

183. Gaiha GD, Dong T, Palaniyar N, Mitchell DA, Reid KB, Clark HW. Surfactant protein A binds to HIV and inhibits direct infection of CD4+ cells, but enhances dendritic cell-mediated viral transfer. J Immunol. 2008;181(1):601–9.

